# Distinct roles for dopamine clearance mechanisms in regulating behavioral flexibility

**DOI:** 10.1101/823401

**Authors:** Clio Korn, Thomas Akam, Kristian HR Jensen, Cristiana Vagnoni, Anna Huber, Elizabeth M Tunbridge, Mark E Walton

## Abstract

Dopamine plays a crucial role in adaptive behavior, and dysfunctional dopamine is implicated in multiple psychiatric conditions characterized by inflexible or inconsistent choices. However, the precise relationship between dopamine and flexible decision making remains unclear. One reason is that, while many studies have focused on the activity of dopamine neurons, efficient dopamine signaling also relies on clearance mechanisms, notably the dopamine transporter (DAT), which predominates in striatum, and catechol-O-methyltransferase (COMT), which predominates in cortex. The exact locus, extent, and timescale of the effects of DAT and COMT are uncertain. Moreover, there is limited data on how acute disruption of either mechanism affects flexible decision making strategies mediated by cortico-striatal networks. To address these issues, we combined pharmacological modulation of DAT and COMT with electrochemistry and behavior in mice. DAT blockade, but not COMT inhibition, regulated sub-second dopamine release in the nucleus accumbens core, but surprisingly neither clearance mechanism affected evoked release in prelimbic cortex. This was not due to a lack of sensitivity, as both amphetamine and atomoxetine changed the kinetics of sub-second release. In a multi-step decision making task where mice had to respond to reversals in either reward probabilities or the choice sequence to reach the goal, DAT blockade selectively impaired, and COMT inhibition improved, performance after reward reversals, but neither manipulation affected the adaptation of choices after action-state transition reversals. Together, our data suggest that DAT and COMT shape specific aspects of behavioral flexibility by regulating striatal and cortical dopamine, respectively, at fast and slow timescales.

## Introduction

Changes in behavioral flexibility are a central component of multiple psychiatric and neurological conditions, such as schizophrenia and substance use disorders, that involve dysfunctional dopamine transmission (1,2). However, the precise relationship between flexible behavior and patterns of dopamine signaling in different brain regions remains a matter of contention. This gap in our knowledge has important clinical implications, since therapeutics for schizophrenia or Parkinson’s disease that target dopamine transmission and successfully remediate some symptoms have been reported to have limited influence on – or even to impair – behavioral flexibility (3).

Rapid and precisely-timed dopamine activity plays a crucial role in adaptive behavior, signaling how much better or worse an event is than predicted (a “reward prediction error”). Boosting or blocking these phasic signals can change the likelihood of repeating or switching away from a recent choice (4). However, effective dopamine signaling depends not just on dopamine neuron activity but also on clearance mechanisms, which are critical for the temporal and spatial regulation of dopamine’s actions. Crucially, clearance mechanisms vary across cortico-striatal networks implicated in flexible decision making (2). Recycling via the dopamine transporter (DAT) predominates in nucleus accumbens (NAc) and other striatal regions (5), while enzymatic degradation, particularly by catechol-*O*-methyltransferase (COMT), is more prominent in prefrontal cortex (PFC) (6,7). These clearance mechanisms are ideally placed to shape mesolimbic and mesocortical dopamine control over flexible decision making. Indeed, recent evidence suggests that manipulations of either the mesolimbic pathway to NAc or mesocortical pathway to medial PFC can have distinct effects on behavioral flexibility (8–10).

While there have been a number of studies investigating how DAT and COMT shape reward pursuit and cognitive functions like working memory, respectively (11–15), their role in regulating behavioral flexibility has been comparatively neglected. Individual variation in DAT or COMT function in humans has been shown to correlate with differences in reinforcement learning and flexible decision making (16–22). However, it is not clear precisely how markers of individual variation in transporter or enzyme function in humans relate to dopamine signaling *in vivo*. Notably, when the effects of targeted pharmacological or genetic manipulations of DAT have been investigated in animal models, deficits in reward learning or behavioral flexibility have been notably absent despite changes in motivation (11,12,23,24), while evidence from animal models of COMT’s involvement in behavioral flexibility is limited (25). Moreover, there is still much uncertainty over the locus, extent, and timescale of the effects of DAT and COMT, particularly concerning their regulation of phasic fluctuations in dopamine that produce reward prediction error-like signals in both striatum and cortex (9,26).

To address these issues, we combined pharmacological modulation of DAT and COMT function with electrochemistry and behavior. We first determined the effects of DAT blockade and COMT inhibition on fast fluctuations in dopamine in NAc using fast-scan cyclic voltammetry (FCV) and extended this approach for the first time to understand the effects of DAT and COMT on stimulated release in PFC. We then studied the effects of each clearance mechanism on behavioral flexibility using a multi-step decision-making task that dissociated changes in the optimal response strategy to reach a goal from changes in reward probabilities at the goal location. Given the evidence that dopamine is predominantly activated by reward prediction errors over sensorimotor surprise (27), we anticipated that any influence of DAT or COMT might be selective to situations where there is a need to update choice strategies following a change in reward value. Together, our data show that DAT and COMT have specific and bidirectional influences over rapid reward-driven behavioral flexibility, but only DAT achieves this via regulation of fast striatal dopamine signaling.

## Methods

See Supplementary Information for detailed methods.

### Animals

For details of animal numbers, see figures and Supplementary Information. Adult male C57BL/6 mice were used because COMT exhibits sexually dimorphic effects (28). Mice were maintained at >85% free feeding weight for all behavioral experiments, except for the sucrose preference test where they were water deprived for 3hrs prior to test sessions. Food and water were otherwise provided *ad libitum*. All experiments were conducted in accordance with the UK Home Office Animals (Scientific Procedures) Act 1986 and the local ethical review board at the University of Oxford.

### Drugs

Mice were administered the selective COMT inhibitor tolcapone (30mg/kg; TRC Inc)(14,29) and/or the selective DAT blocker GBR-12909 dihydrochloride (6mg/kg, Tocris)(30). These doses are effective at altering behavior without causing motor stereotypies (see Supplementary Information). D-amphetamine (in sulfate formulation; Tocris) was used at 4mg/kg (31), and the NET blocker atomoxetine (in hydrochloride formulation; Tocris) at 1mg/kg (32). All drugs were dissolved in 20% hydroxypropyl-beta-cyclodextrin (Acros Organics) in 0.9% saline (AquPharm), which served as a vehicle control in all experiments. All drugs were delivered by intraperitoneal injection.

### Fast-scan cyclic voltammetry (FCV)

Carbon fibre electrodes targeting NAc core or prelimbic PFC were implanted and voltammetric recordings were made as previously described (33). Catecholamine release was induced by electrical stimulation of the ventral tegmental area (VTA) (60 biphasic 2ms pulses, 50Hz, 300μA), based on literature (34,35) and on pilot experiments. Following a 30min pre-drug baseline period, tolcapone or vehicle was administered. GBR-12909 or vehicle was then given after a further 90min of recording. Amphetamine was tested either after the last injection or in naïve animals. Atomoxetine was only administered to drug naïve animals. Data were obtained from 57 mice (n=5-14 per drug group in each brain region, Supplementary Table 1).

### Behavioral tasks

#### Progressive ratio (PR) task

During PR sessions, the number of active lever presses required to obtain each subsequent reward (60uL of 10% sucrose solution) was increased according to the following equation: required lever presses = 5*e^i*0·16^-5 (‘i’: trial number). Drug effects were assessed by giving mice two systemic injections prior to PR test sessions: tolcapone or vehicle (either 105 or 120mins before testing), followed by GBR-12909 or vehicle (15 or 60mins before testing). Each mouse received all possible drug combinations over four PR test sessions according to a counterbalanced within-subjects design.

#### Multi-step decision making task

The task was adapted from the two-step task developed by Daw et al. (36) for dissociating model-based and model-free reinforcement learning in humans, as reported in (37,38). On each trial, mice chose between a top and a bottom nose poke, causing either the left or right reward port to light up, selection of which caused reward (20% sucrose) to be delivered on a probabilistic schedule (Figure 4A). At any point in time, a particular first-step action (top or bottom) usually led to a particular second-step state (left or right port active) (“common” transitions, 80% of trials), though sometimes led to the opposite state (“rare” transitions, 20% of trials). Similarly, one reward port had a high probability of giving reward (0.8) and the other a low reward probability (0.2)(Figure 4B).

Unlike other recent rodent adaptations of multi-step decision tasks (39–41), both the reward probabilities in the second-step states and the transition probabilities linking the first-step actions to the second-step states reversed in blocks (20 trials after an exponential moving average of choices [tau=8 trials] crossed a 75% correct threshold, leading to either a reward or transition reversal).

Once animals had acquired the task, pharmacological manipulations were performed. Tolcapone or its vehicle were administered 90min prior to sessions; GBR-12909 or its vehicle were administered 15min prior to sessions. Each subject received a total of 8 separate sessions of each drug and 5 sessions of each vehicle injection, order counterbalanced across animals.

### Data analysis

Detailed analysis methodologies are reported in Supplementary Information. Full statistical results are presented in figure legends. The datasets are available from the corresponding author on reasonable request.

## Results

### DAT, but not COMT, influences fast dopamine fluctuations in NAc

We first wanted to extend our understanding of how DAT and COMT shape dynamics of catecholamine transmission in striatum and cortex by using FCV to assess the size and timing of release events at sub-second resolution. This is important because the precision of dopamine transients is thought to be critical for appropriate reward learning and behavioral flexibility and because most previous studies have assessed the roles of DAT and COMT on dopamine transmission at relatively slow (minute-level) timescales (42–46).

We initially focused on VTA-evoked release in NAc core. As expected, DAT blockade increased the magnitude and duration of dopamine transients: GBR-12909 increased the peak size of evoked release and slowed both the latency to peak and the rate of decay from peak (GBR-12909 x time interaction: all F>6.5, p<0.021, Figure 1F,H,J). In contrast, COMT inhibition had no effect (all F<2.4, p>0.14) (Figure 1E,G,I). When considered with previous microdialysis studies (42,44,46), this implies that only DAT recycling, and not COMT breakdown, plays a direct role in shaping the dynamics of fast dopamine transmission in NAc.

**Figure 1:**
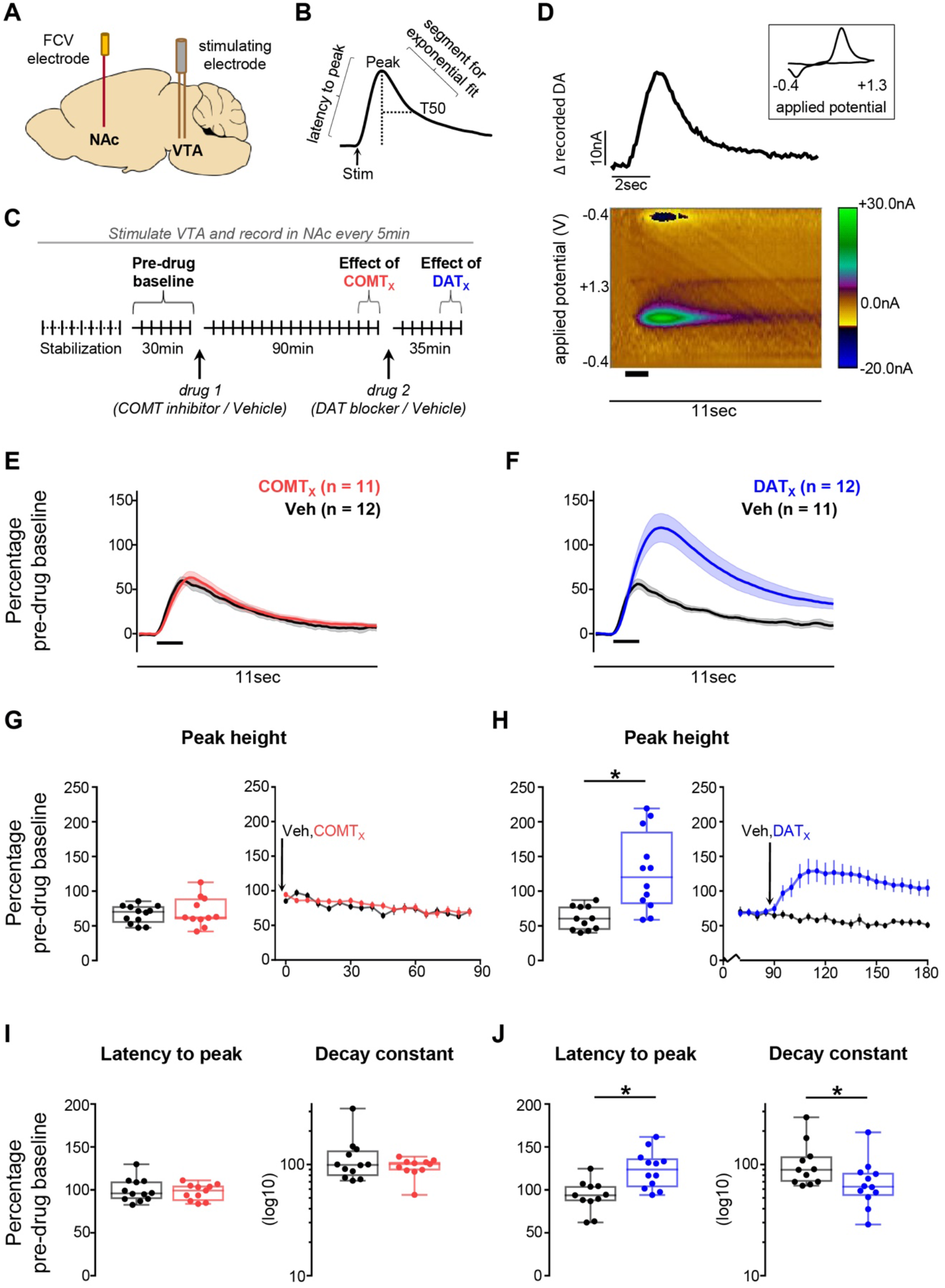
Effects of COMT inhibition and DAT blockade on evoked dopamine release in the NAc. **A)** Schematic of recording and stimulating electrodes in NAc core and VTA respectively (see Supplementary Figure 1 for precise locations). **B)** Illustration of features of dopamine transients quantified during analysis. **C)** Schematic of experiment structure. Drug injections are indicated below the timeline and time points of interest above it. **D)** An example recording made in the NAc core. The pseudocolour plot shows the recorded current as a function of the applied potential of the triangular waveform (y axis) over an 11sec period (x axis); timing and duration of stimulation indicated by thick black bar. The trace above the pseudocolour plot shows the extracted current attributable to dopamine release as a function of time. The cyclic voltammogram (current versus applied voltage; boxed inset) during the peak of the signal, 1.5sec after the time of stimulation, is consistent with dopamine release. **E)** Evoked dopamine release in NAc in animals given the COMT inhibitor as drug 1 (red) compared to release in those given vehicle as drug 1 (black) 85min after the first injection. Release is normalized to the average pre-drug baseline peak height (equivalent to 100% on the y axis), binned over 15min centred on the time point of interest, and presented as mean ± SEM across animals within each drug group. Timing and duration of stimulation indicated by thick black bar. **F)** As in (E), but comparing release in animals given the DAT blocker as drug 2 (blue) with release in those given vehicle as drug 2 (black) 30min after the second injection. **G)** Quantification of the peak height of evoked dopamine release in NAc following administration of the COMT inhibitor. Left: peak height at the same time point shown in (E). Each animal’s data is shown individually and is normalized to its average pre-drug baseline peak height (equivalent to 100% on the y axis). Box plots show median and 25th and 75^th^ percentiles; whiskers extend from the minimum to maximum value. Right: normalized peak height (mean ± SEM for each drug group) over the 90min following the first injection in animals that received the COMT inhibitor compared with those that received vehicle as drug 1. COMT inhibition had no effect on the peak height, latency to peak, or decay from peak of evoked transients. **H)** As in (G), but comparing data from animals that received the DAT blocker with data from those that received vehicle as drug 2; right-hand plot shows peak height over the 90min following the second injection. **I)** As in the left-hand plot of (G), but showing the quantification of the latency from stimulation to peak (left) and of the decay from the peak to T50 (right). Decay constant data are shown on a log_10_ scale for clarity. **J)** As in (I), but for the DAT blocker. **Statistical analysis:** COMT inhibition did not alter the size or kinetics of evoked release (all F<2.4, p>0.14). DAT blockade increased the size of evoked release compared to vehicle (GBR-12909 x time interaction: F_1,19_=18.1, p=0.0004; main effect of GBR-12909: F_1,19_=7.8, p=0.011; main effect of time: F_1,19_=11.5, p=0.003). Post-hoc tests confirmed an increase in evoked release selectively following administration of the DAT blocker but not of vehicle (p=0.002). DAT blockade also slowed the latency to peak compared to vehicle (GBR-12909 x time interaction: F_1,19_=12.6, p=0.002; main effect of GBR-12909: F_1,19_=7.8, p=0.011; main effect of time: F_1,19_=6.3, p=0.021). In addition, there was an interactive effect of the DAT blocker and time on the rate of decay from peak (F_1,19_=6.5, p=0.020). Post-hoc tests confirmed the selective effect of the DAT blocker on the latency to peak (p=0.002) and showed a trend level effect of DAT blockade on postpeak kinetics compared to vehicle (p=0.078), with a difference in the rate of decay in the DAT blockade group before and after receiving the drug (p=0.007) that was not seen in the vehicle group (p=0.565).

### Neither DAT nor COMT affect fast catecholamine fluctuations in PFC

We then examined VTA-evoked release in prelimbic PFC. Transients here were in general considerably smaller than in NAc (pre-drug: NAc 11.48±12.36nA; PFC 2.26±0.96nA, mean±standard deviation: Figure 2B). Unexpectedly, we found little reliable influence of either DAT blockade (Figure 2D,F,H) or COMT inhibition (Figure 2C,E,G) on the size or kinetics of evoked release in PFC (all F<3.7, p>0.07, except GBR-12909 x time interaction F_1,21_=6.0, p=0.023). The lack of observed effect was not because our approach was insufficiently sensitive to detect drug-induced changes in PFC: both amphetamine, an indirect sympathomimetic known to potentiate dopamine (Figure 2I), and atomoxetine, a noradrenaline transporter blocker (Figure S3), changed VTA-evoked release kinetics (all p<0.031). Therefore, although COMT is implicated in shaping minute-by-minute dopamine tone in frontal cortex (25,45,47), our data indicate that neither it nor DAT regulates the release and kinetics of fast catecholamine transmission in this region.

**Figure 2:**
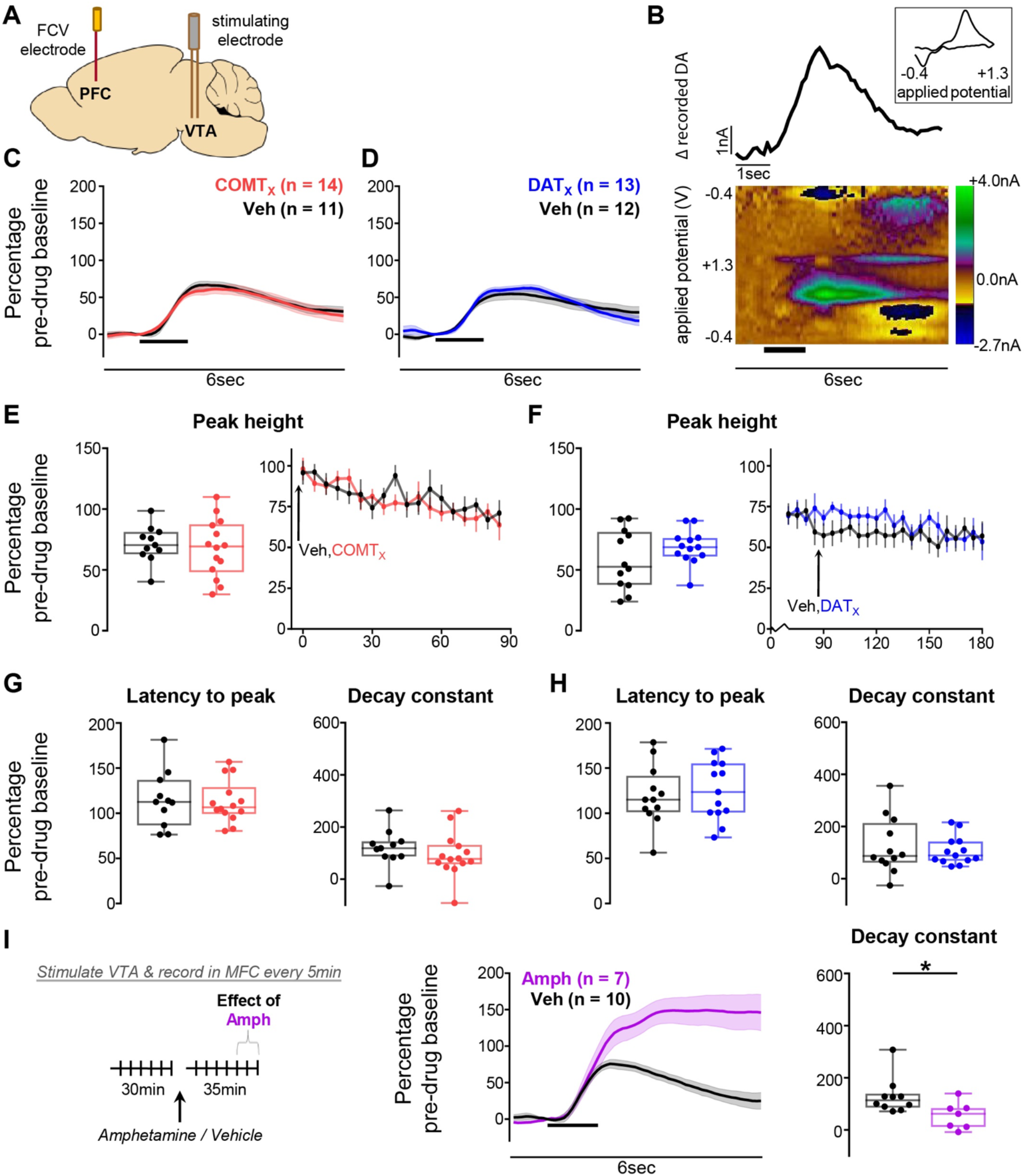
Effects of COMT inhibition and DAT blockade on evoked dopamine release in the PFC. **A)** Schematic of recording and stimulating electrodes in prelimbic cortex and the VTA respectively (see Supplementary Figure 2 for precise histology). **B)** An example recording made in the prelimbic PFC. The pseudocolour plot shows the recorded current as a function of the applied potential of the triangular waveform (y axis) over a 6sec period (x axis); timing and duration of stimulation indicated by thick black bar. The trace above the pseudocolour plot shows the extracted current attributable to dopamine release as a function of time. The cyclic voltammogram (current versus applied voltage; boxed inset) during the peak of the signal, 1.4sec after the time of stimulation, is consistent with catecholamine release. **C)** Evoked dopamine release in the PFC in animals given the COMT inhibitor as drug 1 (red) compared to release in those given vehicle as drug 1 (black) 85min after the first injection. Release is normalized to the average pre-drug baseline peak height (equivalent to 100% on the y axis), binned over 15min centred on the time point of interest, and presented as mean ± SEM across animals within each drug group. Timing and duration of stimulation indicated by thick black bar. **D)** As in (C), but comparing release in animals given the DAT blocker as drug 2 (blue) with release in those given vehicle as drug 2 (black) 30min after the second injection. **E)** Quantification of the peak height of evoked dopamine release in PFC following administration of the COMT inhibitor. Left: peak height at the same time point shown in (C). Each animal’s data is shown individually and is normalized to its average pre-drug baseline peak height (equivalent to 100% on the y axis). Box plots show median and 25th and 75th percentiles; whiskers extend from the minimum to maximum value. Right: normalized peak height (mean ± SEM for each drug group) over the 90min following the first injection in animals that received the COMT inhibitor compared with those that received vehicle as drug 1. **F)** As in (E), but comparing data from animals that received the DAT blocker with data from those that received vehicle as drug 2; right-hand plot shows peak height over the 90min following the second injection. **G)** As in the left-hand plot of (E), but showing the quantification of the latency from stimulation to peak (left) and of the decay from the peak to T50 (right). **H)** As in (G), but for the DAT blocker. **l)** The effect of amphetamine (magenta) on evoked dopamine release in PFC. Left: schematic of experiment structure. Middle: evoked dopamine release normalized to the average pre-drug baseline peak height, binned over 15min centred on the time point of interest (30min after drug injection), and presented as mean ± SEM across animals within each group. Right: quantification of the signal decay, with each animal’s data shown individually and normalized to its average pre-drug baseline decay constant. **Statistical analysis:** Neither COMT inhibition nor DAT blockade altered any measure of evoked release in PFC (all F<3.7, p>0.07, with the exception of a GBR-12909 x time interaction on post-peak signal decay (F_1,21_=6.0, p=0.023), which likely reflects the difficulty of performing an exponential fit on small cortical transients rather than to a drug effect). Amphetamine both increased the peak height (independent samples t-test: t_15_=-4.6, p=0.0003) and decreased the decay constant value (t_15_=2.4, p=0.030) compared to vehicle.

### DAT blockade, but not COMT inhibition, increases motivation to work for reward

We investigated how differential regulation of dopamine clearance by DAT and COMT influences motivation to work for reward. While striatal dopamine and DAT have been linked to motivational drive by numerous previous studies (11-13,48), much less is known about COMT’s influence (42,46).

We therefore first examined the separate and interactive effects of DAT and COMT on motivation to work for reward on a progressive ratio (PR) task. DAT blockade increased motivational drive indexed by willingness to work for reward and sustained engagement. It selectively increased active lever presses (main effect of GBR-12909: F_1,19_=26.4, p<0.001, Figure 3C,D) and speeded re-engagement latencies (main effect of GBR-12909: F_1,19_=15.4, p=0.001, Figure 3E,F). In contrast, there were no reliable effects of COMT inhibition, either in isolation or in interaction with DAT blockade, on any measures in the PR task (all F<3.1, p>0.09, Figure 3C-F).

**Figure 3:**
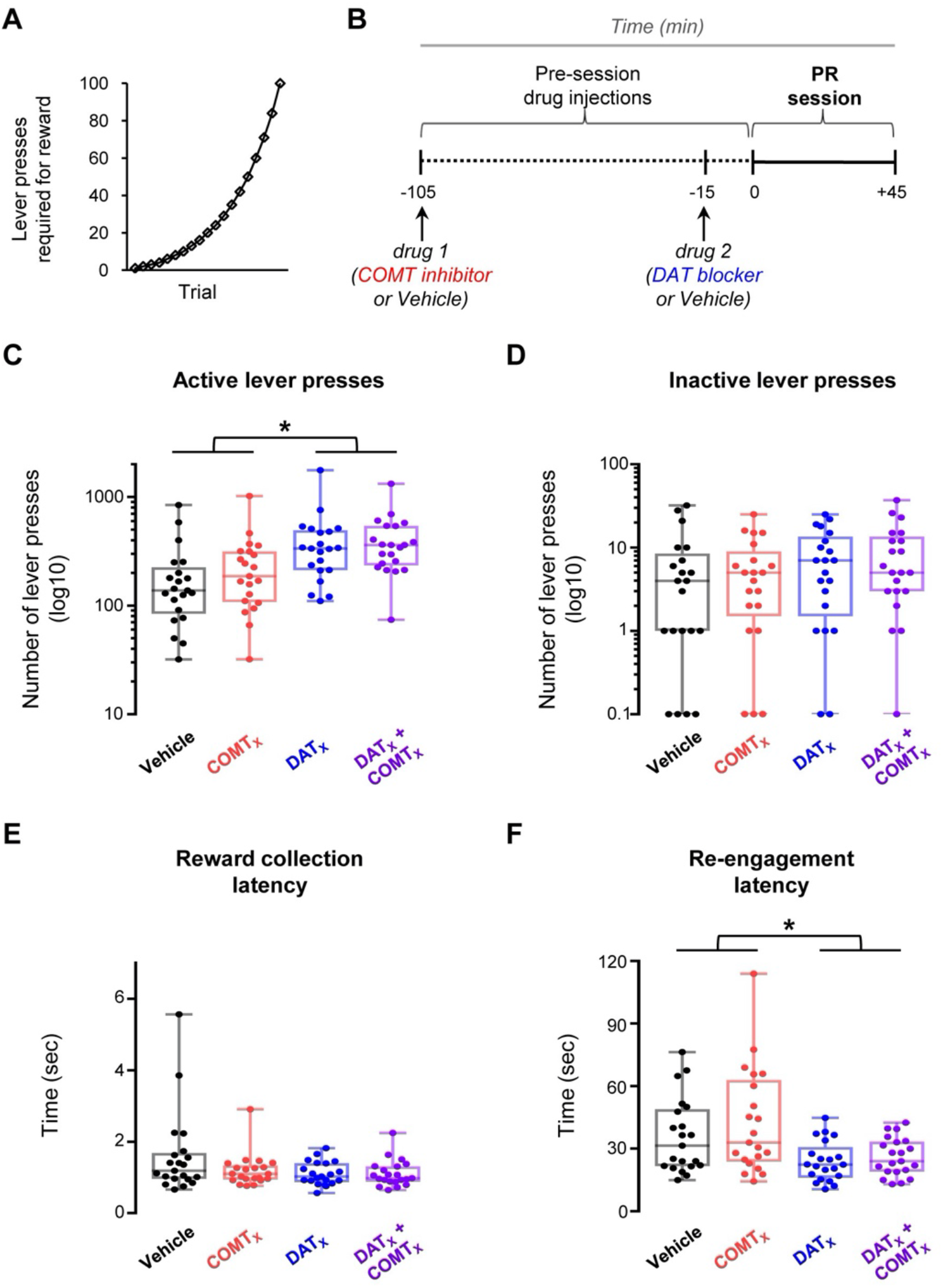
Effects of DAT blockade and COMT inhibition on the progressive ratio (PR) task. A within-subjects design was used to test drug effects; n=21. **A)** Depiction of the progression across trials of lever press ratios required to obtain reward during PR sessions. **B)** Schematic of experiment structure. Drug injections are indicated below the timeline. Timing of drug administration shown is for cohort 2 (see Supplementary Methods). **C)** Number of responses on the active lever during the PR session under the four drug 1 & drug 2 treatment conditions: vehicle & vehicle (black), COMT inhibitor & vehicle (red), vehicle & DAT blocker (blue), and COMT inhibitor & DAT blocker (purple). Each data point shows the lever press session total for one animal. Boxplots show median and 25^th^ and 75^th^ percentiles; whiskers extend from the minimum to maximum value. Lever press data are shown on a log_10_ scale for clarity. **D)** As in (C), but for responses on the inactive lever. Data points again show lever press session totals for individual animals and are displayed on a log_10_ scale. **E)** As in (C), but for the latency to collect reward following its delivery. Each data point shows the cross-trial average latency for one animal. **F)** As in (C), but for the latency to re-engage with the task by recommencing lever pressing following the consumption of reward. Data points show cross-trial average latencies for individual animals. **Statistical analysis:** DAT blockade increased active lever presses (main effect of GBR-12909: F_1,19_=26.4, p=6×10^-5^) and speeded task performance (main effect of GBR-12909 on re-engagement latency: F_1,19_=15.4, p=0.001; main effect of GBR-12909 on reward collection latency: F_1,19_=4.1, p=0.056) but did not affect inactive lever presses (F_1,19_=2.7, p=0.118). COMT inhibition did not alter PR task performance, either on its own or when combined with DAT blockade (all F<3.1, p>0.09).

### DAT and COMT have distinct effects on value updating during multi-step decision making

The results of our voltammetry and PR experiments largely reinforce traditional views of DAT’s and COMT’s roles in dopamine transmission and behavior, emphasizing DAT’s role in shaping striatal dopamine transmission and aspects of motivational drive. However, given the importance of precisely-regulated dopamine transmission for appropriate reward-guided learning and flexible decision making, we hypothesized that DAT and COMT might affect these processes in the context of a more complex decision-making task likely to recruit both striatal and cortical circuitry (Figure 4A,B). Unlike other similar rodent multi-step paradigms (39–41), our task included reversals in both reward *and* action-state transition probabilities. Crucially, this allowed us to examine the influence of DAT and COMT over different aspects of behavioral flexibility: how efficiently mice respond either when the location of high probability reward changes (reward reversals) or when the initial choice required to reach the high probability reward changes (transition reversals).

Mice became proficient at the task, triggering reversals on average in <30 trials prior to drug administration, and performing on average 434-799 trials during each of 26 experimental sessions (Figure 4C). We first examined the influence of DAT or COMT on willingness-to-work given that the actions incurred negligible effort-related cost. Consistent with our PR experiment, DAT blockade but not COMT inhibition increased the rate at which subjects performed trials (veh/drug x DAT/COMT interaction: F_1,7_=6.52, p=0.038, Figure 4D,F). There was also a selective speeding of second-step reaction times following DAT blockade, particularly following rare transitions (veh/drug x DAT/COMT interaction: F_1, 7_=7.76, p=0.027, veh/drug x common/rare interaction: F_1,7_=9.26, p=0.019, Figure 4E,G). These findings further support a selective role for DAT, but not COMT, in invigorating reward-guided responding.

**Figure 4:**
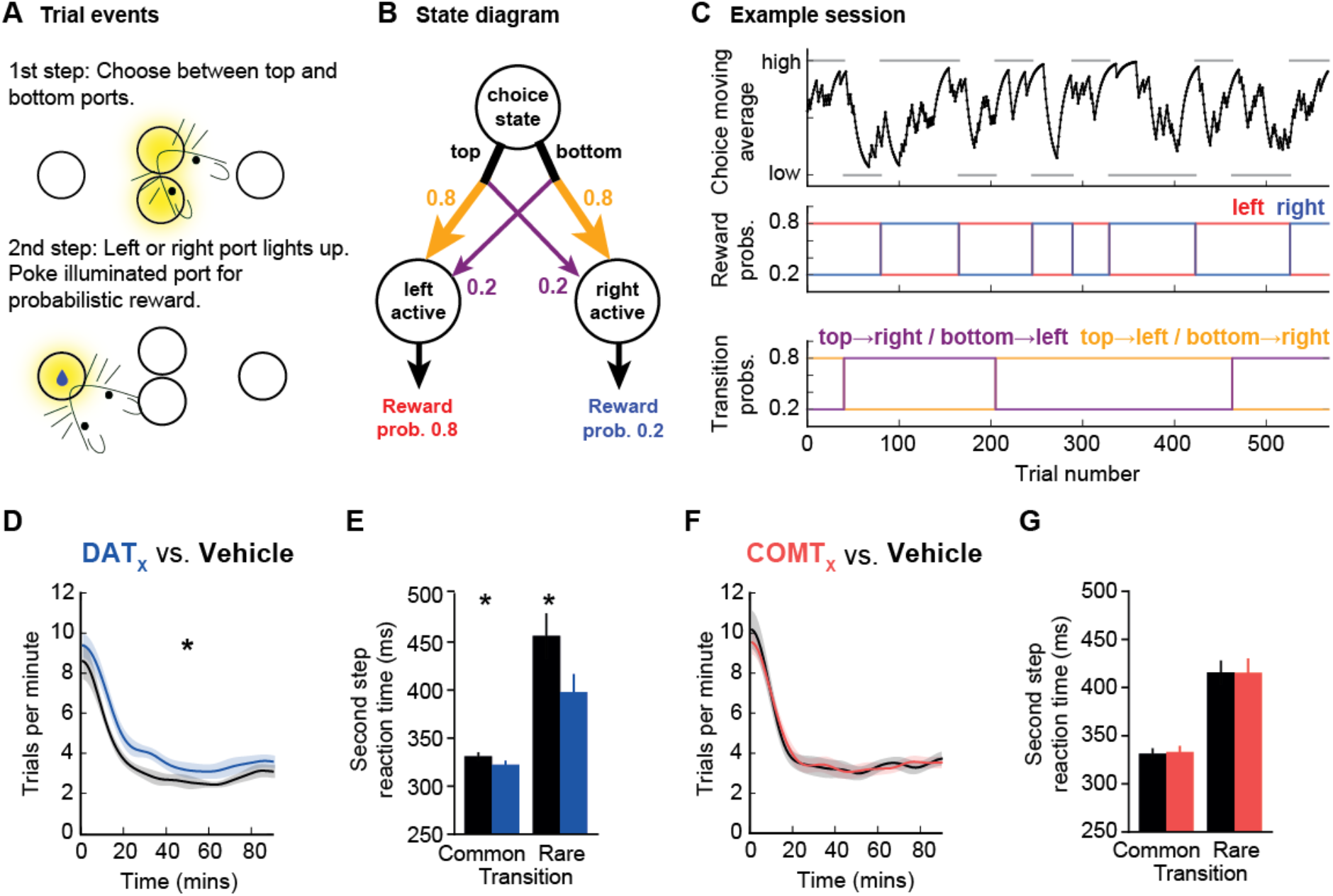
Effects of DAT blockade and COMT inhibition on performance during a multi-step decision making task. A within-subjects design was used to test drug effects; n=8, 26 vehicle/drug sessions per mouse (8 x GBR-12909, 8 x tolcapone, 5 x each drug’s vehicle), average range = 434-799 trials per session per mouse. **A)** Diagram of two-step task apparatus and trial events. **B)** Diagram of the task state space, reward and transition probabilities are shown for one block type **C)** Example session. Top panel: black line shows exponential moving average of subjects’ choices (tau = 8 trials). The correct choice (high or low) for each block is indicated by the grey bars. Middle panel: reward probabilities for each block for the left (red) and right (blue) sides. Bottom panel: transition probabilities linking first step actions to second step states, yellow line shows probability of top→left and bottom→right transitions, purple line shows probability of top→right and bottom→left transitions. Colours in middle and bottom panel of (C) match those in (B). **D,F)** Trial rate as a function of session time (D, DAT blockade; F, COMT inhibition). Trial rates were smoothed with a Gaussian of 5 minute standard deviation. Shaded areas show cross-subject SEM. **E,G)** Median second-step reaction times following common and rare transitions (E, DAT blockade; G, COMT inhibition). The second-step reaction time is the time from choosing high or low to entering the active side port at the second step. Error bars show cross subject SEM. **Statistical analysis:** DAT blockade, but not COMT inhibition, increased trial rate (main effect of veh/drug: F_1,7_=9.79, p=0.016; veh/drug x DAT/COMT interaction: F_1,7_=6.52, p=0.038; post-hoc paired t-tests, DAT: p=0.012, COMT: p=0.98). DAT blockade, but not COMT inhibition, also speeded second-step reaction times (main effect of veh/drug: F_1,7_=8.20, p=0.024; veh/drug x DAT/COMT interaction: F_1,7_=7.76, p=0.027), particularly following rare transitions (veh/drug x common/rare interaction: F_1,7_=9.26, p=0.019; 3-way interaction did not quite reach significance: veh/drug x common/rare x DAT/COMT interaction: F_1,7_=4.35, p=0.076). *, p<0.05.

We next assessed how DAT blockade and COMT inhibition affected choice performance, focusing on reward and transition reversals. Neither drug affected pre-reversal choice rates at the end of blocks (all F_1,7_<1.59, p>0.24). Following reward reversals, DAT blockade significantly slowed (p=0.0012) and COMT inhibition significant improved (p=0.0496) adaptation, as measured by the choice probability trajectories on and off drug (Figure 5A,C). Importantly, however, neither drug affected the speed of adaptation following reversals in transition probabilities (all p>0.22) (Figure 5B,D; Supplementary Table 3,4). To further examine the specificity of this reversal effect, we directly compared whether choices immediately after each type of reversal were selectively influenced by drug administration. This again highlighted that the effect of DAT blockade and COMT inhibition depended on whether the reward or transition probabilities reversed (veh/drug x DAT/COMT x reward/transition-reversal interaction: F_1,7_=6.88, p=0.034). Specifically, the drugs selectively influenced adaptation after reversals in the reward probabilities (veh/drug x DAT/COMT interaction: F_1,7_=17.85, p=0.004) but had no influence after reversals in transition probabilities (all F<0.36, p>0.56). Taken together, these data demonstrate that dopamine clearance mechanisms shape adaptive behaviors only when there is a change in reward location and not when the action sequence required to reach that reward reverses.

**figure 5:**
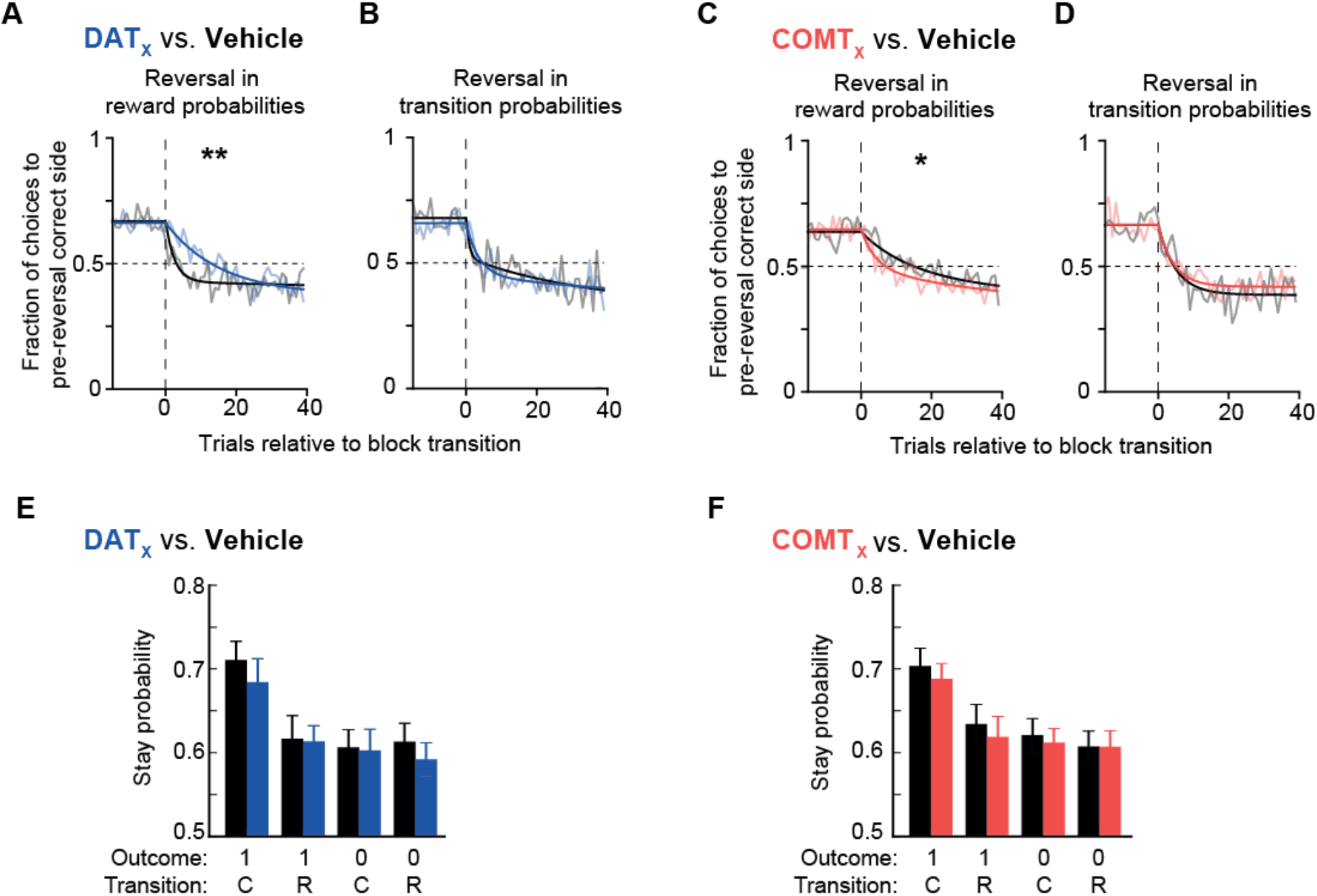
DAT blockade impairs and COMT inhibition improves value updating, but not transition updating, during multi-step decision making. **A-D)** Choice probability trajectories around reversals in reward probabilities (A, DAT blockade, C, COMT inhibition) and transition probabilities (B, DAT blockade, D, COMT inhibition). Pale lines show cross-subject mean choice probability trajectory and dark lines show double exponential fit. **E,F)** Probability of repeating the first step choice on the next trial as a function of the trial outcome (rewarded or not) and experience state transition (common or rare) (E, DAT blockade, F, COMT inhibition). **Statistical analysis:** Neither DAT blockade nor COMT inhibition changed the choice rates at the end of each block prior to a reversal (all F<1.59, p>0.24). DAT blockade and COMT inhibition both modulated the speed of reward reversals, assessed by fitting a double exponential to the choice probability trajectory and using permutation testing to evaluate whether differences between drug and corresponding vehicle condition were significant (reward reversals: GBR-12909: p=0.0012; tolcapone: p=0.0496; transition reversals: all p>0.23). In addition, direct comparison of post-reversal choices highlighted that the effect depended on whether the reward or transition probabilities reversed (veh/drug x DAT/COMT x reward/transition-reversal interaction: F_1,7_=6.88, p=0.034; reversals in the reward probabilities: veh/drug x DAT/COMT interaction: F_1,7_=17.85, p=0.004; reversals in transition probabilities: all F<0.36, p>0.56). A mixed effects logistic regression showed that the mice’s choices were sensitive to both the outcome (reward/no reward) *and* the transition structure (common/rare), both p<0.001). However, neither of these were reliably affected by drug condition (p>0.086 for all interactions of outcome and transition with drug condition). Instead, the drug manipulations – and particularly DAT blockade – affected the regression model’s ‘correct’ predictor, which captures a general tendency to repeat choices commonly associated with the high reward probability option (correct x veh/drug x DAT/COMT interaction: p=0.016; DAT blockade, p=0.056; COMT inhibition: p=0.28). *, p<0.05; **, p<0.01

To gain insight into what might be causing these changes in behavioral flexibility, we investigated the specific influence of DAT and COMT on decision making strategies and trial-to-trial learning. Consistent with previous work using the same task (38), mice were sensitive to the transition structure of the task, not simply being more likely to repeat any choice that was rewarded, as would be expected if they were exclusively using a ‘model-free’ reinforcement learning strategy, but instead being more likely to repeat a rewarded choice after a common than a rare transition (Figure 5E,F). A comparison of reinforcement learning models showed that mice’s choices were best explained by a model that employed both model-free and model-based control (see Supplementary Information for discussion of the mice’s strategies, Figure S4). Moreover, a logistic regression analysis of the factors influencing choice repetition demonstrated a significant effect of both outcome and transition (p<0.001 mixed effects logistic regression).

Importantly, and more unexpectedly, neither DAT nor COMT manipulations reliably modified these measures, as might be expected if either mechanism were simply disrupting trial-by-trial reward learning (p>0.086 for all interactions of outcome and transition with drug condition). There was also no effect on choice or motor biases (p>0.21). Instead, the drug manipulations - and particularly DAT blockade - affected the regression model’s ‘correct’ predictor, which captures a general tendency to repeat choices commonly associated with the high reward probability option (correct x veh/drug x DAT/COMT interaction: p=0.016). Together, these results imply that DAT blockade impairs reward-guided behavioral flexibility not by biasing decision making strategies or through a direct effect on reward learning, but instead by disrupting the cumulative effect of past choices and outcomes on reward-driven choices.

## Discussion

Here we demonstrate distinct roles for DAT and COMT in flexible reward-guided decision making. FCV recordings confirmed a role for DAT recycling, but not COMT degradation, in regulating fast fluctuations in NAc dopamine transmission but showed that neither affected evoked transients in PFC, indicating that clearance mechanisms other than DAT and COMT contribute to regulation of cortical dopamine at sub-second timescales. Behaviorally, we confirmed that DAT blockade, but not COMT inhibition, increased motivational drive and promoted task engagement, as previously observed (11,12). Importantly, we additionally demonstrated specific and bidirectional roles for dopamine clearance mechanisms in behavioral flexibility, with DAT impairing and COMT causing a relative improvement in performance selectively following reversals in reward probabilities.

Previous studies have indicated that DAT can affect motivation but has limited, if any, influence over reward learning (11-13,18,23,48). Here, we demonstrated that DAT not only modulates motivational performance and task engagement but, notably, can also influence behavioral flexibility. Specifically, DAT blockade selectively disrupted the ability to adapt following reversals in reward probabilities. Strikingly, the effect of DAT blockade was not observed following reversals in the transition probabilities linking first-step choices to second-step reward states, even though these also required animals to adapt their behavior (in this case, at the first-step choice) in order to maximize rewards obtained. Therefore, the effect cannot be attributed to a general behavioral inflexibility, a finding also supported by the lack of influence on any motor level biases in the logistic regression model. Instead, our data are consistent with DAT influencing rapid reward-driven alternations in behavioral policies in addition to shaping motivational components of reward-guided behavior.

In agreement with previous studies (42,44,46), we found that DAT blockade both increased and extended evoked NAc dopamine transients. The effects were similar to those seen on spontaneous transients in freely-moving animals following administration of a catecholamine transporter blocker or stimulant drugs (31,49). In addition, we found no effect of DAT blockade on evoked release in prelimbic PFC, consistent with the sparse DAT expression in this region (50). Therefore, DAT blockade predominantly exerts its effects through regulation of striatal dopamine.

Given that fast fluctuations in striatal dopamine correlate with reward prediction error signals, which are strongly linked to animals’ ability to form certain reward-related associations (27,51), it may seem surprising that there is such limited evidence in animal models linking DAT to regulation of adaptive behavior. However, although the reward reversal impairment might initially seem to imply a change in reward learning, there was in fact no reliable evidence that DAT blockade caused alterations to specific trial-by-trial reinforcement learning or a change in reinforcement learning strategies. Instead, it appears that DAT more prominently helps to regulate how the history of past choices and outcomes influences behavioral policies, which can shape how quickly animals detect a change in reward value. Such a conclusion aligns with studies showing effects of dopamine on flexible responding that go beyond influences on trial-by-trial reinforcement learning. For instance, DAT knockdown mice, which have increased tonic dopamine though, unlike here, reduced phasic transients, have reduced flexibility in response to changes in effort cost that has been linked to a reduced sensitivity to recent outcome history (52). Similarly, a human DAT polymorphism specifically influenced perseverative errors during a probabilistic reward reversal task (19). These findings imply that the behavioral consequences of DAT blockade on reward learning and behavioral flexibility will depend strongly on how changeable and uncertain rewards are in any context.

Research on COMT’s role in behavior has largely focused on cognitive functions; its significance for reward-guided behavior remains relatively unstudied (22). We found that COMT inhibition had no effect on motivational aspects of behavior, consistent with its lack of effect on NAc dopamine (42,53,54). More surprisingly, COMT inhibition had no effect on the size or kinetics of VTA-evoked transients in prelimbic PFC. This was not due to a lack of sensitivity to sub-second changes in cortex, as amphetamine increased VTA-evoked transient size and slowed the decay of cortical signals, while the norephinephrine transporter blocker atomoxetine slowed signal decay in PFC. Taken together with previous studies, this pinpoints COMT’s selective influence in regulating cortical dopamine transmission over longer timescales than are required for precise reinforcement learning (25,45,47). These results, however, contrast with a report of differences in dopamine overflow in COMT knockout mice (34). While this discrepancy may reflect different analytical approaches, it may also speak to differences between the acute changes in COMT in adult animals induced by inhibition, compared to the constitutive changes present in the knockout mouse, in which compensatory changes are possible (14). This highlights the need for a greater understanding of the timescales over which compensatory changes across cortical and subcortical dopamine systems occur (55–57).

Although COMT does not appear to regulate the kinetics of shorter-lived dopamine transients, COMT inhibition nonetheless selectively speeded adaptation to reward reversals relative to vehicle injections. The precise mechanism by which COMT regulates reward-driven behavioral flexibility remains unclear, as there were no consistent changes in behavioral strategy or reinforcement learning that emerged from the analyses. However, although acute pharmacological inhibition is different than the chronic changes in enzymatic activity arising from the human COMT Val/Met polymorphism (58), our findings are concordant with reports of increased sensitivity to rewards and faster reinforcement learning in Met allele carriers, who have lower COMT activity than Val allele homozygotes (59,60).

Together, our data suggest that DAT and COMT differentially regulate specific aspects of reward-based behavioral flexibility. Surprisingly, however, we found no evidence that either does this by influencing the balance of reinforcement learning strategies or shaping trial-by-trial learning. These findings demonstrate the importance of different regulators of dopamine signaling, operating over distinct timescales and in different brain regions, in shaping multiple aspects of adaptive reward-guided behavior beyond reinforcement learning. Such data therefore highlight the complexities associated with the development of novel pharmacological agents to remedy the dopamine-dependent changes seen in patients with psychiatric disorders.

## Acknowledgements

We thank Greg Daubney for performing histology; Richard Keithley for advice and assistance with chemometric analysis; and Katie Jennings, Paul Harrison, and Jaime McCutcheon for useful discussions about the data. This research was funded in whole, or in part, by the Wellcome Trust (4-year studentship WT096586 to CK, fellowships WT090051MA and 202831/Z/16/Z to MEW), as well as the Royal Society (EMT) and the BBSRC (project grant: BB/M024148/1). For the purpose of open access, the author has applied a CC BY public copyright licence to any Author Accepted Manuscript version arising from this submission.

## Author Contributions

CK, EMT and MEW conceived the project and, with TA, designed the experiments. CK, KHJ, CV, and AH collected the data. CK and TA analyzed the data, and CK, TA, EMT and MEW wrote the manuscript.

## References

1. Izquierdo A, Jentsch JD. Reversal learning as a measure of impulsive and compulsive behavior in addictions. Psychopharmacology (Berl). 2012 Jan 1;219(2):607–20.

2. Izquierdo A, Brigman JL, Radke AK, Rudebeck PH, Holmes A. The neural basis of reversal learning: An updated perspective. Neuroscience. 2017 Mar 14;345:12–26.

3. Swainson R, Rogers RD, Sahakian BJ, Summers BA, Polkey CE, Robbins TW. Probabilistic learning and reversal deficits in patients with Parkinson’s disease or frontal or temporal lobe lesions: possible adverse effects of dopaminergic medication. Neuropsychologia. 2000 May 1;38(5):596–612.

4. Hamid AA, Pettibone JR, Mabrouk OS, Hetrick VL, Schmidt R, Vander Weele CM, et al. Mesolimbic dopamine signals the value of work. Nat Neurosci. 2016 Jan;19(1):117–26.

5. Sulzer D, Cragg SJ, Rice ME. Striatal dopamine neurotransmission: Regulation of release and uptake. Basal Ganglia. 2016 Aug;6(3):123–48.

6. Karoum F, Chrapusta SJ, Egan MF. 3-Methoxytyramine Is the Major Metabolite of Released Dopamine in the Rat Frontal Cortex: Reassessment of the Effects of Antipsychotics on the Dynamics of Dopamine Release and Metabolism in the Frontal Cortex, Nucleus Accumbens, and Striatum by a Simple Two Pool Model. J Neurochem. 1994;63(3):972–9.

7. Tunbridge EM, Harrison PJ, Weinberger DR. Catechol-o-Methyltransferase, Cognition, and Psychosis: Val158Met and Beyond. Biol Psychiatry. 2006 Jul;60(2):141–51.

8. Radke AK, Kocharian A, Covey DP, Lovinger DM, Cheer JF, Mateo Y, et al. Contributions of nucleus accumbens dopamine to cognitive flexibility. Eur J Neurosci. 2019 Aug;50(3):2023–35.

9. Ellwood IT, Patel T, Wadia V, Lee AT, Liptak AT, Bender KJ, et al. Tonic or Phasic Stimulation of Dopaminergic Projections to Prefrontal Cortex Causes Mice to Maintain or Deviate from Previously Learned Behavioral Strategies. J Neurosci. 2017 Aug 30;37(35):8315–29.

10. Verharen JPH, de Jong JW, Roelofs TJM, Huffels CFM, van Zessen R, Luijendijk MCM, et al. A neuronal mechanism underlying decision-making deficits during hyperdopaminergic states. Nat Commun. 2018 Feb 21;9(1):731.

11. Cagniard B, Balsam PD, Brunner D, Zhuang X. Mice with chronically elevated dopamine exhibit enhanced motivation, but not learning, for a food reward. Neuropsychopharmacology. 2006;31 (7):1362–70.

12. Cagniard B, Beeler JA, Britt JP, McGehee DS, Marinelli M, Zhuang X. Dopamine Scales Performance in the Absence of New Learning. Neuron. 2006 Sep;51(5):541–7.

13. Peciña S, Cagniard B, Berridge KC, Aldridge JW, Zhuang X. Hyperdopaminergic mutant mice have higher “wanting” but not “liking” for sweet rewards. J Neurosci. 2003;23(28):9395–402.

14. Barkus C, Korn C, Stumpenhorst K, Laatikainen LM, Ballard D, Lee S, et al. Genotype-Dependent Effects of COMT Inhibition on Cognitive Function in a Highly Specific, Novel Mouse Model of Altered COMT Activity. Neuropsychopharmacology. 2016 Aug 10;41:3060–9.

15. Farrell SM, Tunbridge EM, Braeutigam S, Harrison PJ. COMT Val158Met Genotype Determines the Direction of Cognitive Effects Produced by Catechol-O-Methyltransferase Inhibition. Biol Psychiatry. 2012 Mar;71(6):538–44.

16. Colzato LS, Waszak F, Nieuwenhuis S, Posthuma D, Hommel B. The flexible mind is associated with the catechol-O-methyltransferase (COMT) Val158Met polymorphism: Evidence for a role of dopamine in the control of task-switching. Neuropsychologia. 2010 Jul;48(9):2764–8.

17. Aarts E, Roelofs A, Franke B, Rijpkema M, Fernández G, Helmich RC, et al. Striatal Dopamine Mediates the Interface between Motivational and Cognitive Control in Humans: Evidence from Genetic Imaging. Neuropsychopharmacology. 2010 Aug;35(9):1943–51.

18. Kaiser RH, Treadway MT, Wooten DW, Kumar P, Goer F, Murray L, et al. Frontostriatal and Dopamine Markers of Individual Differences in Reinforcement Learning: A Multi-modal Investigation. Cereb Cortex. 2018 Dec 1;28(12):4281–90.

19. den Ouden HEM, Daw ND, Fernandez G, Elshout JA, Rijpkema M, Hoogman M, et al. Dissociable Effects of Dopamine and Serotonin on Reversal Learning. Neuron. 2013 Nov 20;80(4):1090–100.

20. Krugel LK, Biele G, Mohr PNC, Li S-C, Heekeren HR. Genetic variation in dopaminergic neuromodulation influences the ability to rapidly and flexibly adapt decisions. Proc Natl Acad Sci U S A. 2009 Oct 20;106(42):17951–6.

21. Doll BB, Bath KG, Daw ND, Frank MJ. Variability in Dopamine Genes Dissociates Model-Based and Model-Free Reinforcement Learning. J Neurosci. 2016 Jan 27;36(4):1211–22.

22. Tunbridge EM, Huber A, M Farrell S, Stumpenhorst K, J Harrison P, E Walton M. The role of catechol-O-methyltransferase in reward processing and addiction. CNS Neurol Disord-Drug Targets Former Curr Drug Targets-CNS Neurol Disord. 2012;11(3):306–23.

23. Costa VD, Tran VL, Turchi J, Averbeck BB. Dopamine modulates novelty seeking behavior during decision making. Behav Neurosci. 2014;128(5):556–66.

24. Seu E, Lang A, Rivera RJ, Jentsch JD. Inhibition of the norepinephrine transporter improves behavioral flexibility in rats and monkeys. Psychopharmacology (Berl). 2009 Jan 1;202(1):505–19.

25. Tunbridge EM, Bannerman DM, Sharp T, Harrison PJ. Catechol-O-Methyltransferase Inhibition Improves Set-Shifting Performance and Elevates Stimulated Dopamine Release in the Rat Prefrontal Cortex. J Neurosci. 2004 Jun 9;24(23):5331–5.

26. Hart AS, Rutledge RB, Glimcher PW, Phillips PEM. Phasic Dopamine Release in the Rat Nucleus Accumbens Symmetrically Encodes a Reward Prediction Error Term. J Neurosci. 2014 Jan 15;34(3):698–704.

27. Schultz W. Dopamine reward prediction-error signalling: a two-component response. Nat Rev Neurosci. 2016 Mar;17(3):183–95.

28. Harrison PJ, Tunbridge EM. Catechol-O-methyltransferase (COMT): a gene contributing to sex differences in brain function, and to sexual dimorphism in the predisposition to psychiatric disorders. Neuropsychopharmacology. 2008;33(13):3037–45.

29. Männistö PT, Kaakkola S. Catechol-O-methyltransferase (COMT): Biochemistry, Molecular Biology, Pharmacology, and Clinical Efficacy of the New Selective COMT Inhibitors. Pharmacol Rev. 1999 Dec 1;51(4):593–628.

30. Izenwasser S, Werling LL, Cox BM. Comparison of the effects of cocaine and other inhibitors of dopamine uptake in rat striatum, nucleus accumbens, olfactory tubercle, and medial prefrontal cortex. Brain Res. 1990;520(1-2):303–9.

31. Daberkow DP, Brown HD, Bunner KD, Kraniotis SA, Doellman MA, Ragozzino ME, et al. Amphetamine Paradoxically Augments Exocytotic Dopamine Release and Phasic Dopamine Signals. J Neurosci. 2013 Jan 9;33(2):452–63.

32. Bymaster FP, Katner JS, Nelson DL, Hemrick-Luecke SK, Threlkeld PG, Heiligenstein JH, et al. Atomoxetine increases extracellular levels of norepinephrine and dopamine in prefrontal cortex of rat: a potential mechanism for efficacy in attention deficit/hyperactivity disorder. Neuropsychopharmacology. 2002;27(5):699–711.

33. Syed ECJ, Grima LL, Magill PJ, Bogacz R, Brown P, Walton ME. Action initiation shapes mesolimbic dopamine encoding of future rewards. Nat Neurosci. 2016 Jan;19(1):34–6.

34. Yavich L, Forsberg MM, Karayiorgou M, Gogos JA, Mannisto PT. Site-Specific Role of Catechol-O-Methyltransferase in Dopamine Overflow within Prefrontal Cortex and Dorsal Striatum. J Neurosci. 2007 Sep 19;27(38):10196–209.

35. Yorgason JT, Jones SR, España RA. Low and high affinity dopamine transporter inhibitors block dopamine uptake within 5 sec of intravenous injection. Neuroscience. 2011 May;182:125–32.

36. Daw ND, Gershman SJ, Seymour B, Dayan P, Dolan RJ. Model-Based Influences on Humans’ Choices and Striatal Prediction Errors. Neuron. 2011 Mar;69(6):1204–15.

37. Akam T, Costa R, Dayan P. Simple Plans or Sophisticated Habits? State, Transition and Learning Interactions in the Two-Step Task. PLOS Comput Biol. 2015 Dec 11;11(12):e1004648.

38. Akam T, Rodrigues-Vaz I, Marcelo I, Zhang X, Pereira M, Oliveira RF, et al. The Anterior Cingulate Cortex Predicts Future States to Mediate Model-Based Action Selection. Neuron. 2021 Jan 6;109:1–15.

39. Groman SM, Massi B, Mathias SR, Curry DW, Lee D, Taylor JR. Neurochemical and behavioral dissections of decision-making in a rodent multi-stage task. J Neurosci. 2018 Nov 9;2219–18.

40. Hasz BM, Redish AD. Deliberation and Procedural Automation on a Two-Step Task for Rats. Front Integr Neurosci. 2018 Aug 3;12:30.

41. Miller KJ, Botvinick MM, Brody CD. Dorsal hippocampus contributes to model-based planning. Nat Neurosci. 2017 Sep;20(9):1269–76.

42. Budygin EA, Gainetdinov RR, Kilpatrick MR, Rayevsky KS, Männistö PT, Wightman RM. Effect of tolcapone, a catechol-O-methyltransferase inhibitor, on striatal dopaminergic transmission during blockade of dopamine uptake. Eur J Pharmacol. 1999;370(2):125–31.

43. Carboni E, Silvagni A, Vacca C, Di Chiara G. Cumulative effect of norepinephrine and dopamine carrier blockade on extracellular dopamine increase in the nucleus accumbens shell, bed nucleus of stria terminalis and prefrontal cortex. J Neurochem. 2006 Jan 1;96(2):473–81.

44. Huotari M, Gainetdinov R, Männistö PT. Microdialysis Studies on the Action of Tolcapone on Pharmacologically-Elevated Extracellular Dopamine Levels in Conscious Rats. Pharmacol Toxicol. 1999;85(s1):233–8.

45. Lapish CC, Ahn S, Evangelista LM, So K, Seamans JK, Phillips AG. Tolcapone enhances food-evoked dopamine efflux and executive memory processes mediated by the rat prefrontal cortex. Psychopharmacology (Berl). 2009 Jan;202(1-3):521–30.

46. Raevskii KS, Gainetdinov RR, Budygin EA, Mannisto P, Wightman M. Dopaminergic transmission in the rat striatum in vivo in conditions of pharmacological modulation. Neurosci Behav Physiol. 2002;32(2):183–8.

47. Käenmäki M, Tammimäki A, Myöhänen T, Pakarinen K, Amberg C, Karayiorgou M, et al. Quantitative role of COMT in dopamine clearance in the prefrontal cortex of freely moving mice: Quantitative role of COMT in the prefrontal cortex. J Neurochem. 2010 Sep;114(6):1745–55.

48. Yin HH, Zhuang X, Balleine BW. Instrumental learning in hyperdopaminergic mice. Neurobiol Learn Mem. 2006 May;85(3):283–8.

49. Robinson DL, Wightman RM. Nomifensine amplifies subsecond dopamine signals in the ventral striatum of freely-moving rats. J Neurochem. 2004;90(4):894–903.

50. Sesack SR, Hawrylak VA, Matus C, Guido MA, Levey AI. Dopamine axon varicosities in the prelimbic division of the rat prefrontal cortex exhibit sparse immunoreactivity for the dopamine transporter. J Neurosci. 1998;18(7):2697–708.

51. Flagel SB, Clark JJ, Robinson TE, Mayo L, Czuj A, Willuhn I, et al. A selective role for dopamine in stimulus-reward learning. Nature. 2011 Jan 6;469(7328):53–7.

52. Beeler JA, Daw N, Frazier CRM, Zhuang X. Tonic Dopamine Modulates Exploitation of Reward Learning. Front Behav Neurosci. 2010 Nov 4;4:170.

53. Acquas E, Carboni E, Ree RHA, Prada M, Chiara G. Extracellular Concentrations of Dopamine and Metabolites in the Rat Caudate After Oral Administration of a Novel Catechol-O-Methyltransferase Inhibitor Ro 40-7592. J Neurochem. 1992;59(1):326–30.

54. Garris PA, Wightman RM. Distinct pharmacological regulation of evoked dopamine efflux in the amygdala and striatum of the rat in vivo. Synapse. 1995;20(3):269–79.

55. Meyer-Lindenberg A, Kohn PD, Kolachana B, Kippenhan S, McInerney-Leo A, Nussbaum R, et al. Midbrain dopamine and prefrontal function in humans: interaction and modulation by COMT genotype. Nat Neurosci. 2005 May;8(5):594–6.

56. Pycock CJ, Carter CJ, Kerwin RW. Effect of 6-Hydroxydopamine Lesions of the Medial Prefrontal Cortex on Neurotransmitter Systems in Subcortical Sites in the Rat. J Neurochem. 1980 Jan 1;34(1):91–9.

57. Thompson TL, Moss RL. In vivo stimulated dopamine release in the nucleus accumbens: modulation by the prefrontal cortex. Brain Res. 1995 Jul 17;686(1):93–8.

58. Tunbridge EM, Narajos M, Harrison CH, Beresford C, Cipriani A, Harrison PJ. Which Dopamine Polymorphisms Are Functional? Systematic Review and Meta-analysis of COMT, DAT, DBH, DDC, DRD1-5, MAOA, MAOB, TH, VMAT1, and VMAT2. Biol Psychiatry. 2019 Oct 15;86(8):608–20.

59. Corral-Frías NS, Pizzagalli DA, Carré JM, Michalski LJ, Nikolova YS, Perlis RH, et al. COMT Val158Met genotype is associated with reward learning: a replication study and meta-analysis. Genes Brain Behav. 2016;15(5):503–13.

60. Lancaster TM, Heerey EA, Mantripragada K, Linden DEJ. Replication study implicates COMT val158met polymorphism as a modulator of probabilistic reward learning. Genes Brain Behav. 2015 Jul;14(6):486–92.

