## Supplementary Information for "Distinct roles for dopamine clearance mechanisms in regulating behavioral flexibility"

### Supplementary Methods

#### Animals

Male C57BL/6 mice were obtained from Envigo (formerly Harlan). Mice were aged 9-26 weeks for behavioral experiments and 10-16 weeks for voltammetry experiments. Animals were housed on a 12/12-hr light/dark cycle; all behavioral tests were conducted during the light phase. Mice were food deprived to 85-90% of free feeding weight for the locomotor activity, progressive ratio, and multi-step decision making tasks, and were water deprived for 3hrs prior to test sessions for the sucrose preference test. Food and water were provided *ad libitum* in all other cases. All mice were habituated to handling – including the restraint position used during injections – before experiments began. Care and testing of all animals was conducted under the auspices of the UK Home Office laws and guidelines for the treatment of animals under scientific procedures and of the local ethical review board at the University of Oxford.

#### Drugs

Mice were administered the selective, brain-penetrant COMT inhibitor tolcapone (TRC Inc) and/or the selective DAT blocker GBR-12909 dihydrochloride (Tocris). Tolcapone was used at a 30mg/kg dose, which previous studies have shown affects both dopaminergic transmission and behavior in rodents (1–5). GBR-12909 dihydrochloride was administered at 6mg/kg, a dose that in pilot experiments produced increased locomotor activity but not stereotypy, as assessed using criteria adapted from (6) for use in mice (data not shown). D-amphetamine (in sulfate formulation; Tocris) was used at 4mg/kg, and the NET blocker atomoxetine (in hydrochloride formulation; Tocris) at 1mg/kg. All drugs were dissolved in 20%

hydroxypropyl-beta-cyclodextrin (Acros Organics) in 0.9% saline (AquPharm), which served as a vehicle control in all experiments. All drugs were delivered by intraperitoneal injection, with an injection volume of either 5mL/kg or 10mL/kg (multi-step decision making task only). Drug administration timings were designed to account for differences in drug time courses of action.

Fast-scan cyclic voltammetry (FCV)

A total of 100 mice were used for FCV recordings, including for method development, piloting of electrode placements, and optimization of dopamine recordings. Of these, 14 animals did not provide analyzable data, due to failed drug injections (NAc n = 6; PFC n = 4) or death prior to the completion of the experiment (NAc n = 3; PFC n = 1). For those animals that did provide a complete data set, dopamine signals were required to meet a set of quality control criteria: (1) a ratio of standard deviation of pre-stimulation noise to peak height ≤ 0.5; (2) a detectable peak consistently occurring ≤ 3sec after stimulation; and (3) peak timing varying by ≤ 2sec across 30min bins. For the kinetics analysis, the fit of the exponential decay function was required to have $R^2 > 0.5$. A final cohort of 57 mice with electrodes located in the NAc or PFC contributed to the dataset reported in this study. Group sizes for each region and drug treatment of the between-subjects design are shown in Supplementary Table 1.

*Electrode fabrication and implantation*

Recording and reference electrodes were made in-house (7,8) and pre-calibrated in a flow cell (9) to allow conversion of recorded signals from current (nA) into concentration (nM). When calibration factors were unavailable (9 electrodes), a mean calibration factor was used. The stimulating electrodes were untwisted, bipolar, stainless steel electrodes measuring 0.15mm in diameter (PlasticsOne).

Mice were implanted with electrodes under anesthesia (isoflurane induction [Abbott Laboratories Ltd] and urethane maintenance [Sigma Aldrich]). Mice were administered glycopyrronium bromide (0.01-0.02mg/kg i.p. alongside urethane; MercuryPharm Ltd) and glucosesaline (0.5% in 0.9% saline; ~7mL every 3hrs; Aquapharm), and temperature was maintained at 35.0-37.0°C with a rectal probe and heating blanket. Recording electrodes were placed either in the NAc core (+1.40 anterior-posterior [AP] and -0.75 medial-lateral [ML] from bregma; -3.50 to -4.75 dorsal-ventral [DV] from the skull) or prelimbic PFC (+2.50 AP and -0.50 ML from bregma, -1.30 to -2.30 DV from skull). Stimulating electrodes targeted the posterior-medial portion of the VTA, where the cell bodies of dopamine neurons projecting to both NAc and PFC are located (-3.50 AP and -0.35 ML from bregma; -4.00 to -4.80 DV from brain surface) (10–12) . A reference electrode was placed in the contralateral hemisphere (+4.80 AP and +1.00 ML from bregma).

*FCV recordings*

Voltammetric recordings were made as previously described (8). Transient release was induced by passing current through the stimulating electrode targeting the VTA via a DS3 Stimulator (Digitimer) (stimulation parameters: pulse number = 60 pulses, frequency = 50Hz, amplitude = 300μA, pulse width = 2ms, pulse phase = biphasic), based on previous literature (13,14) and on pilot experiments that established the parameters required to reliably evoke detectable neurotransmitter release in both the NAc and PFC, where the lower density of dopamine terminals reduces signal-to-noise compared to striatum. Stimuli were generated and recordings collected using Tarheel CV (National Instruments).

Evoked signals decayed over time and so were allowed to stabilize (stimulating every 5-10mins for ~2.5hrs during NAc, and ~1hr during PFC, recordings). After the stabilization period, stimulations were made every 5min. Following a 30min pre-drug baseline period, tolcapone or vehicle was administered. GBR-12909 or vehicle was then given after a further

90min of recording, and recordings continued for several hours. Amphetamine was tested in two groups of animals: those that had received tolcapone or vehicle and naïve animals. There were no differences in signal decay between these groups (data not shown), so they were combined. Atomoxetine was only administered to drug naïve animals. Data were averaged into 15min bins for analysis: the effects of tolcapone were assessed at 85mins, of GBR-12909 and amphetamine at 30mins, and of atomoxetine at 60min post-administration. Once recording was complete, electrode placement was ascertained as previously described (8).

Behavioral tasks

*Sucrose preference test*

Sucrose preference was assessed in open-top cages equipped with two water bottles. 12 mice (previously run on locomotor/stereotypy test for drug dosage assessment and counterbalanced for prior drug exposure) were tested for 5hrs each day for a total of 7 days (Days 1-3: water exposure only; Days 4-7: sucrose exposure). Bottles were weighed immediately before and after testing to determine consumption. On day 7, mice received two injections – the first (tolcapone or vehicle) 1hr before testing and the second (GBR-12909 or vehicle) immediately before testing. Preference for sucrose solution (10% weight/volume; Sigma Aldrich) was assessed as a ratio of sucrose consumption to total consumption.

*Progressive ratio task*

A total of 24 mice (two cohorts: n = 12 previously used for locomotor activity and sucrose preference tests, counterbalanced for prior drug exposure; n = 12 test naïve, given a vehicle injection 3 days prior to testing) were tested on the progressive ratio task. Of these, 3 mice were excluded from the analysis: 2 due to failed injections and 1 that was unable to learn the complete task. The timing of experimental drug administration differed between the two cohorts. Cohort 1 received the first injection 120min and the second injection 60min before the start of the session, whereas cohort 2 received the first injection 105min and the second injection 15min before the session. No notable effects of cohort were observed (data not shown), so findings from the two cohorts are reported together.

The task was conducted as previously described (15) in standard operant chambers (Med Associates Inc). Rewards consisted of 60μL of 10% sucrose solution. Animals were trained on increasing fixed ratio (FR) schedules until they were able to earn ≥ 15 rewards and achieved an active:inactive lever press ratio of ≥ 3:1 on an FR5 schedule over 2 consecutive days. During progressive ratio (PR) sessions, the number of active lever presses required to obtain each subsequent reward was increased according to the following equation: number of required lever presses = $5*e^{i*0.16} - 5$ ('i': trial number). Drug effects on behavior were assessed by giving mice two systemic injections prior to PR test sessions: tolcapone or vehicle, followed by GBR-12909 or vehicle. Each mouse received all possible drug combinations over four PR test sessions according to a counterbalanced within-subjects design. Drug testing days were interleaved with two washout days during which no injections were given: one day of testing on an FR5 schedule and one day of testing on a PR schedule.

*Multi-step decision making task*

The task was adapted from the two-step task developed by (16) for dissociating model-based and model-free reinforcement learning in humans, as reported in (17,18). We had capacity to test 8 animals on this task. In order to optimize our training process, we began preliminary training with a total of 16 mice for 6 days. The 8 animals that had performed the most trials during that time then continued on to complete an additional 32 days of training and subsequent drug testing. Note that as all drug testing was done on a within-subjects basis, there is no possibility of this initial selection biasing the results.

The task was run in 8 custom made 12x12cm operant boxes controlled using pyControl (https://pycontrol.readthedocs.io). The behavioral apparatus consisted of 4 nose poke ports; a 'top' and a 'bottom' port in the center flanked by 'left' and 'right' ports (Figure 4A). Each trial started with the top and bottom ports lighting up. The subject chose top or bottom, causing either the left or right port to light up. The subject then poked the illuminated side for a probabilistic reward (20% weight/volume sucrose solution; Sigma Aldrich). At any point in time, one reward port had a high probability of giving reward (0.8) and the other a low reward probability (0.2). Similarly, a particular first-step action (top or bottom) usually led to a particular second-step state (left or right port active) ("common" transitions, 80% of trials), though sometimes led to the opposite state ("rare" transitions, 20% of trials) (Figure 4B). Both the reward probabilities in the second-step states and the transition probabilities linking the first-step actions to the second-step states reversed in blocks. Block transitions were triggered based on the subject's behavior, occurring 20 trials after an exponential moving average (tau = 8 trials) of choices crossed a 75% correct threshold (Figure 4C). Reversals in reward probability occurred twice as often as reversals in transition probability. The two types of reversals did not occur simultaneously.

Subjects encountered the full trial structure from the first day of training. The only task parameters that were changed over the course of training were the state and reward transition probabilities and the reward sizes; reward size was gradually reduced and the reward and transition probabilities gradually adjusted over 38 days of training as mice became progressively more engaged with the task and learned to perform it better (see Supplementary Table 2 for details). All animals had at least 12 sessions with the final task parameters prior to drug administration. Animals were considered fully trained and ready for pharmacological testing when the group average met the following criteria: (1) a 3-day average of >400 trials per session, (2) a 3-day average of >4 reversals per session, and (3) a 3-day average combined reversal learning speed of <30 trials. By the end of training, animals were performing 442.2±28.8 trials and 5.7±0.6 blocks per session, completing 70.5±5.2 trials per block, and obtaining reward on 51.6±0.5 percent of trials (mean±SEM across animals over the 3 days before the first injection).

Pharmacological manipulations were performed every second day of testing using a within-subjects design. On intervening days mice were run on the task but received no injection. Tolcapone or its vehicle control were administered 90min prior to the start of sessions; GBR-12909 or its vehicle control were administered 15min prior to the start of sessions. Subjects received a total of 8 of each drug and 5 of each vehicle injection, with order counterbalanced across animals.

Experimental design and statistical analyses

Statistical analyses were conducted in SPSS versions 20 and 24 (IBM Computing), with the exception of the multi-step decision-making task (described further below), with significance set at $\alpha = 0.05$. With the exceptions noted below, data were analyzed using analysis of variance (ANOVA), with drug group(s) (and time, where relevant) as factors. For repeated-measures ANOVAs, Greenhouse-Geisser corrections were applied where data failed Mauchley's test of sphericity. Simple main effects analyses were conducted as necessary when significant interactions were found. Least square difference pairwise comparison tests were used to assess which groups were driving any significant main or interactive effects.

*FCV recording analysis*

FCV recordings were processed using software written in LabVIEW and custom Matlab scripts. Dopamine levels were extracted using a chemometric approach based on training sets from individual animals (19–21). Cyclic voltammograms were low-pass filtered at 2kHz and background subtracted using the 5 scans prior to stimulation.

Drug effects on evoked dopamine were assessed by quantifying several features of the evoked transients, including: the peak height; the latency from the start of stimulation to the peak; and the rate of decay of the signal from the peak to T50 (the time when the signal had decayed to half the peak height) (Figure 1B). (In cases where the signal did not fall to half its peak height, the decay over the 3sec following the peak was used.) Prior to statistical analysis, parameters were normalized to the pre-drug baseline signal and binned across three individual recordings at the time of interest. The significance of drug effects on each parameter was determined using repeated-measures ANOVAs, with time as the within-subjects factor and drug 1 (tolcapone or vehicle) and drug 2 (GBR-12909 or vehicle) as between-subjects factors. As we found no interactive effects of the two experimental drugs, the effects of COMT inhibition and DAT blockade are presented separately.

Note that as the cortex is innervated by significant noradrenergic as well as dopaminergic fibers (22,23), and because dopamine and noradrenaline have very similar cyclic voltammograms (24,25), signals recorded in cortex using FCV can only be identified as catecholaminergic, not as definitively dopaminergic. However, previous studies have shown that electrical stimulation of the VTA predominantly evokes dopamine rather than noradrenaline release in PFC (26) and there are also several examples of optical and electrical stimulation of VTA dopamine neurons causing increases in dopamine levels in cortex (27,28). The VTA placement used here causes substantial increases in dopamine release in NAc. It has also been shown that electrical stimulation of mouse VTA causes dopamine-dependent changes in frontal cortex activity (29). Finally, the noradrenergic pathways from the LC terminate most strongly in superficial layers of medial frontal cortex, while the majority of our recordings likely came from deeper cortical layers. Therefore, it is highly likely that the recorded PFC transients are principally dopaminergic, even though a contribution of noradrenaline cannot be entirely ruled out.

*Progressive ratio task analysis*

The main outcome measure was cumulative active lever presses over the session. We also examined cumulative inactive lever presses as a measure of general activity levels, the average reward collection latency, and the average re-engagement latency (the interval between the animal exiting the magazine after reward delivery and its next lever press). The significance of drug effects on each of these measures was assessed using a repeated-measures ANOVA, with drug 1 (tolcapone or vehicle) and drug 2 (GBR-12909 or vehicle) as within-subjects factors.

*Multi-step decision making task analysis*

For the multi-step decision making task, analysis of pharmacological manipulations was restricted to the first 90 minutes of each session. Except where stated otherwise, drug effects were evaluated using repeated-measures ANOVAs. As the two different drugs each had a corresponding vehicle condition (see above), within-subjects factors were *vehicle/drug* – differentiating both vehicle from both drug conditions – and *GBR-12909/tolcapone* – differentiating GBR-12909 and its respective vehicle from tolcapone and its respective vehicle. Drug effects on trial rate (Figure 4D,F) were quantified as the mean number of trials each subject performed over the first 90 minutes. Drug effects on second-step reaction times (Figure 4E,G) were quantified using the median reaction time for each subject for each condition, with common/rare transition as an additional within-subjects factor.

Drug effects on subjects' ability to track which option had highest reward probability were quantified by looking at the fraction of correct choices at the end of blocks and the speed of adaptation to reversals (Figure 5A-D). The fraction of correct choices in the last 15 trials of each block was evaluated for each subject in drug and vehicle conditions and compared using a repeated-measures ANOVA. Two different approaches were used to quantify the speed of adapting to reversals. The fraction of correct choices in the first 15 trials of each block was calculated for each subject separately following reversals in the reward and transition probabilities and compared using a repeated-measures ANOVA with reversal type (reward or transition) and drug condition as within subject factors. To get a more fine-grained picture of how adaptation to reversals was affected by the drugs, the choice probability trajectory following reversals was fit by a sum of two exponential decays defined by the equation:

$$P_n = P_c - (P_c - P_0)(w_f e^{-\frac{n}{\tau_f}} + (1 - w_f)e^{-\frac{n}{\tau_s}})$$

Where $P_n$ is the probability on trial $n$ of choosing the option that was correct following the reversal; $P_c$ is the asymptotic probability of choosing the correct option, defined as the cross-subject mean fraction of correct choices over the last 15 trials of all blocks; $P_0$ is the initial probability of choosing the correct option (calculated as $1 - P_c^{\prime}$, where $P_c^{\prime}$ is the fraction of correct choices at the end of blocks preceding reversals of type being analyzed); $\tau_f$ is the time constant of the fast exponential decay; $\tau_s$ is the time constant of the slow exponential decay; and $w_f$ is the weighting applied to the fast component relative to the slow component.

We used permutation testing to evaluate whether differences between drug and corresponding vehicle condition were significant, with the analysis performed independently for GBR-12909 and tolcapone. The curve was fit using a squared error cost function to the cross-subject mean choice probability trajectory for drug and vehicle conditions, and the difference $\Delta x_{true}$ between drug and vehicle conditions was evaluated for parameters $\tau_f$, $\tau_s$, $w_f$. We then constructed an ensemble of 5000 permuted datasets in which the assignments of sessions to the drug and vehicle conditions were randomized. Randomization was performed within subjects, such that the number of sessions from each subject in each condition was preserved. For each permuted dataset we re-ran the analysis and evaluated the difference in each parameter between the two conditions, to give a distribution of $\Delta x_{perm}$, which in the limit of many permutations is the distribution of $\Delta x$ under the null hypothesis that there is no difference between the conditions. The two-tailed p value for the observed difference is given by:

$$\mathcal{P} = 2 \min\left(\frac{\mathrm{M}}{\mathcal{N}}, 1 - \frac{\mathrm{M}}{\mathcal{N}}\right)$$

Where $\mathcal{N}$ is the number of permutations and $\mathcal{M}$ is the number of permutations for which $\Delta x_{perm} > \Delta x_{true}$.

The statistical significance of trial-to-trial learning, and its modulation by drug treatment, was assessed using a logistic regression model. The model predicted repeating the previous choice as a function of trial outcome (rewarded or not), transition (common or rare), and their interaction. We additionally included two predictors capturing choice biases: one for a bias towards the top / bottom poke, and one for rotational bias – i.e. a tendency to choose top / bottom following trials that ended in the left / right second-step, which is observed in some animals on this task (18). We further included a predictor that promotes repeating correct choices. This prevents correlation between action values at the start of the trial and subsequent trial events from biasing on the transition-outcome interaction predictor loading (17). An in depth characterization of subjects behavior on this task (17,18), indicates that model-based reinforcement learning tends to generate a main effect of transition rather than a transition-outcome interaction due to online learning about the transition probabilities.

The regression model was fit to a dataset comprising tolcapone, GBR-12909, and their corresponding vehicle sessions. Drug effects were modeled by interacting each predictor with *vehicle/drug* – differentiating both vehicle from both drug conditions – and *GBR-12909/tolcapone* – differentiating GBR-12909 and its respective vehicle from tolcapone. All coefficients were treated as independent random effects across subjects and the resulting hierarchical regression was fit using the lme4 mixed effects package in the R statistical language (R Development Core Team, 2010). Random effects whose variance fit to zero were removed from the model to enable the fit to converge. This did not remove random effects for any predictors with significant fixed effects. P values were calculated using the

LmerTest package (30) using Satterthwaite's method for approximating degrees of freedom. The regression model fit to the complete dataset indicated significant three-way interactions between model predictors, vehicle/drug and GBR-12909/tolcapone. To unpack what was driving this interaction, we subsequently performed separate model fits for each drug with its corresponding vehicle, interacting the base predictors with vehicle/drug condition.

We also explored whether fitting reinforcement learning (RL) models to the multi-step decision making task data could provide insight into the behavioral strategies used by the mice and how the drugs affected these. To do this, we compared a set of models generated by adding or removing single features from the model found to best describe behavior on this task in (18) (see below for RL model equations). These features included: forgetting about the values and state transitions for not-chosen actions, action perseveration effects spanning multiple trials, and representation of actions both at the level of the choice they represent (e.g. top poke) and the motor action they require (e.g. left → top movement) (for full details see (18)). The reinforcement learning model fitting used a Bayesian hierarchical modelling framework (18,31) in which parameters for individual sessions were drawn from Gaussian distributions at the population level, with the mean and variance of the Gaussians fit to maximize data likelihood. Models were compared using the integrated Bayes Information Criterion (BIC) score. We first fit RL models using data from baseline (no drug or vehicle injections) sessions.

In addition to modeling baseline session data, we compared the signed difference between maximum likelihood parameter estimates after administration of GBR-12909 or tolcapone with their respective vehicles (Supplementary Figure 4B). Statistical significance of drug effects was assessed using a permutation test that permuted drug and corresponding vehicle sessions within subjects to assess the distribution of differences between drug and vehicle sessions expected for each parameter under the null hypothesis that the drug and vehicle had no differential effects.

*Multi-step decision making task RL model equations*

The best fitting RL model used in Supplementary Figure 4B used the parameters and variables shown in Supplementary Table 5.

First-step model-free action values were updated as:

$$Q_{mf}(c) \leftarrow (1-\alpha_Q)Q_{mf}(\mathrm{c}) + \alpha_Q V(s) \tag{1}$$

Second-step state values were updated as:

$$V(\mathrm{s}) \leftarrow (1-\alpha_Q)\mathrm{V}(\mathrm{s}) + \alpha_Q r \tag{2}$$

Value forgetting modified the value of actions not chosen and states not visited as:

$$Q_{mf}(c') \leftarrow (1-f_Q)Q_{mf}(\mathrm{c}') \tag{3}$$

$$V(s') \leftarrow (1-f_Q)\mathrm{V}(\mathrm{s}') \tag{4}$$

Action-state transition probabilities used by the model-based system were updated as:

$$P(s|\mathrm{c}) \leftarrow (1-\alpha_T)P(s|\mathrm{c}) + \alpha_T \tag{5}$$

$$P(s'|\mathrm{c}) \leftarrow (1-\alpha_T)P(s'|\mathrm{c}) \tag{6}$$

Transition probability forgetting was implemented as:

$$P(s|\mathrm{c}') \leftarrow (1-f_T)P(\mathrm{s}|\mathrm{c}') + 0.5 f_T \tag{7}$$

$$P(s'|\mathrm{c}') \leftarrow (1-f_T)P(\mathrm{s}'|\mathrm{c}') + 0.5 f_T \tag{8}$$

At the start of each trial, model-based first step action values were calculated as:

$$Q_{mb}(\mathrm{c}) = \sum_s P(s|c)V(s) \tag{9}$$

In addition to learning values for first step choices (e.g. top), model-free values were also learnt for first-step motor actions (e.g. left→top), updated as:

$$Q_{mo}(c,\ s_{t-1}) \leftarrow (1-\alpha_Q)Q_{mo}(c,\ s_{t-1}) + \alpha_Q(\lambda r + (1-\lambda)V(s)) \tag{10}$$

Motor-level model-free value forgetting was implemented as:

$$Q_{mo}(m') \leftarrow (1-f_Q)Q_{mo}(m') \tag{11}$$

Where $m'$ are all motor actions not taken.

Choice perseveration was modelled using a choice history variable $\bar{c}$ that was an exponential moving average of recent choices, updated as:

$$\bar{c} \leftarrow (1 - \alpha_c)\bar{c} + \alpha_c(c - 0.5) \tag{13}$$

where $c = 1$ if choice is top and $c = 0$ if choice is bottom.

In addition to choice perseveration, the model included motor-level perseveration, modelled using variables $\bar{m}(s_{t-1})$ which were exponential moving averages of choices following trials ending in state $s_{t-1}$, updated as:

$$\bar{m}(s_{t-1}) \leftarrow (1 - \alpha_m)\bar{m}(s_{t-1}) + \alpha_m(c - 0.5) \tag{14}$$

Net action values were given by a weighted sum of model-free, motor-level model-free and model-based action values, biases and perseveration.

$$Q_{net}(c) = G_{mf}Q_{mf}(\text{c}) + G_{mo}Q_{mo}(c, s_{t-1}) + G_{mb}Q_{mb}(\text{c}) + X(c) \tag{15}$$

Where $G_{mf}$, $G_{mo}$ and $G_{mb}$ are weights controlling the influence of respectively the model-free, motor-level model-free and model-based action values, and $X(c)$ is biases and perseveration where:

$$X(top) = B_c + B_r(s_{t-1} - 0.5) + P_c\bar{c} + P_m\bar{m} \tag{16}$$

$$X(bottom) = 0 \tag{17}$$

where $s_{t-1}$ = 1 if previous second step state is left and 0 if right.

Net action values determined choice probabilities via the softmax decision rule:

$$P(c) = \frac{e^{Q_{net}(c)}}{\sum_c e^{Q_{net}(c)}} \tag{18}$$

### Supplementary Results

<u>Sucrose preference test</u>

Prior to starting operant testing, we assessed the effects of DAT blockade and COMT inhibition on basic reward processing using a sucrose preference test. Neither DAT blockade nor COMT inhibition affected either choice of sucrose solution over water or total fluid consumption (all $F<1.7$, $p>0.21$, univariate ANOVA) (data not shown).

Reinforcement learning models of multi-step decision making

In line with previous reports (18), behavior in the baseline sessions was best explained by fitting a model that employed a mixture of model-based and model-free control, along with several additional factors that captured features of the animals' behavior such as choice and motor perseveration, motor biases, forgetting of non-chosen values, and state-action transitions, (Supplementary Figure 4A). The only difference between the model found to best fit data on this task in (18) and that in the current study was that in (18) the best fitting model used eligibility traces as part of its model free update (although the parameter that controlled the influence of these fit to a low value) while the best fitting model in the current study did not use eligibility traces.

We next investigated the effects of drug or vehicle injections on parameter estimates of the models. Although this permutation test indicated that some parameters differed between drug and vehicle conditions at either weakly significant or trend level (Supplementary Table 6), the high parameter count of the model (13 parameters) meant that none of the differences survived multiple comparison correction for the number of model parameters.

Nonetheless, it is potentially worth examining whether the RL model parameters affected at an uncorrected significance level by the drugs can provide additional insight into the significant changes observed in the reversal analysis. For instance, the RL model fit argues against an interpretation of the slower adaptation to reversals under DAT blockade as being due to increased perseveration, as the RL model's choice perseveration parameter was actually reduced by DAT blockade ($p = 0.027$). By contrast, COMT inhibition reduced the influence of model-free RL operating at the level of motor actions (e.g. left→top) ($P = 0.028$), which could potentially accelerate adaptation to reversals, as learning at the motor level is inefficient on this task because the true value of the top or bottom port is independent of the motor action taken to select it.

### Supplementary References

1. Barkus C, Korn C, Stumpenhorst K, Laatikainen LM, Ballard D, Lee S, et al. Genotype-Dependent Effects of COMT Inhibition on Cognitive Function in a Highly Specific, Novel Mouse Model of Altered COMT Activity. Neuropsychopharmacology. 2016 Aug 10;41:3060–9.

2. Budygin EA, Gainetdinov RR, Kilpatrick MR, Rayevsky KS, Männistö PT, Wightman RM. Effect of tolcapone, a catechol-O-methyltransferase inhibitor, on striatal dopaminergic transmission during blockade of dopamine uptake. Eur J Pharmacol. 1999;370(2):125–31.

3. Huotari M, Gainetdinov R, Männistö PT. Microdialysis Studies on the Action of Tolcapone on Pharmacologically-Elevated Extracellular Dopamine Levels in Conscious Rats. Pharmacol Toxicol. 1999;85(s1):233–8.

4. Lapish CC, Ahn S, Evangelista LM, So K, Seamans JK, Phillips AG. Tolcapone enhances food-evoked dopamine efflux and executive memory processes mediated by the rat prefrontal cortex. Psychopharmacology (Berl). 2009 Jan;202(1–3):521–30.

5. Tunbridge EM, Bannerman DM, Sharp T, Harrison PJ. Catechol-O-Methyltransferase Inhibition Improves Set-Shifting Performance and Elevates Stimulated Dopamine Release in the Rat Prefrontal Cortex. J Neurosci. 2004 Jun 9;24(23):5331–5.

6. Creese I, Iversen SD. The role of forebrain dopamine systems in amphetamine induced stereotyped behavior in the rat. Psychopharmacologia. 1974;39(4):345–57.

7. Papageorgiou GK, Baudonnat M, Cucca F, Walton ME. Mesolimbic Dopamine Encodes Prediction Errors in a State-Dependent Manner. Cell Rep. 2016 Apr 12;15(2):221–8.

8. Syed ECJ, Grima LL, Magill PJ, Bogacz R, Brown P, Walton ME. Action initiation shapes mesolimbic dopamine encoding of future rewards. Nat Neurosci. 2016 Jan;19(1):34–6.

9. Sinkala E, McCutcheon JE, Schuck MJ, Schmidt E, Roitman MF, Eddington DT. Electrode calibration with a microfluidic flow cell for fast-scan cyclic voltammetry. Lab Chip. 2012;12(13):2403.

10. Lammel S, Hetzel A, Häckel O, Jones I, Liss B, Roeper J. Unique Properties of Mesoprefrontal Neurons within a Dual Mesocorticolimbic Dopamine System. Neuron. 2008 Mar;57(5):760–73.

11. Lammel S, Ion DI, Roeper J, Malenka RC. Projection-Specific Modulation of Dopamine Neuron Synapses by Aversive and Rewarding Stimuli. Neuron. 2011 Jun;70(5):855–62.

12. Lammel S, Lim BK, Malenka RC. Reward and aversion in a heterogeneous midbrain dopamine system. Neuropharmacology. 2014 Jan;76:351–9.

13. Yavich L, Forsberg MM, Karayiorgou M, Gogos JA, Mannisto PT. Site-Specific Role of Catechol-O-Methyltransferase in Dopamine Overflow within Prefrontal Cortex and Dorsal Striatum. J Neurosci. 2007 Sep 19;27(38):10196–209.

14. Yorgason JT, Jones SR, España RA. Low and high affinity dopamine transporter inhibitors block dopamine uptake within 5 sec of intravenous injection. Neuroscience. 2011 May;182:125–32.

15. Sharma S, Hryhorczuk C, Fulton S. Progressive-ratio Responding for Palatable High-fat and High-sugar Food in Mice. J Vis Exp. 2012 May 3;63:e3754.

16. Daw ND, Gershman SJ, Seymour B, Dayan P, Dolan RJ. Model-Based Influences on Humans' Choices and Striatal Prediction Errors. Neuron. 2011 Mar;69(6):1204–15.

17. Akam T, Costa R, Dayan P. Simple Plans or Sophisticated Habits? State, Transition and Learning Interactions in the Two-Step Task. PLOS Comput Biol. 2015 Dec 11;11(12):e1004648.

18. Akam T, Rodrigues-Vaz I, Marcelo I, Zhang X, Pereira M, Oliveira RF, et al. The Anterior Cingulate Cortex Predicts Future States to Mediate Model-Based Action Selection. Neuron. 2021 Jan 6;109:1–15.

19. Heien ML, Khan AS, Ariansen JL, Cheer JF, Phillips PE, Wassum KM, et al. Real-time measurement of dopamine fluctuations after cocaine in the brain of behaving rats. Proc Natl Acad Sci U S A. 2005;102(29):10023–8.

20. Keithley RB, Mark Wightman R, Heien ML. Multivariate concentration determination using principal component regression with residual analysis. TrAC Trends Anal Chem. 2009 Oct;28(9):1127–36.

21. Keithley RB, Carelli RM, Wightman RM. Rank Estimation and the Multivariate Analysis of in Vivo Fast-Scan Cyclic Voltammetric Data. Anal Chem. 2010 Jul;82(13):5541–51.

22. Lindvall O, Björklund A, Divac I. Organization of catecholamine neurons projecting to the frontal cortex in the rat. Brain Res. 1978 Feb 17;142(1):1–24.

23. Slopsema JS, Van Der Gugten J, De Bruin JPC. Regional concentrations of noradrenaline and dopamine in the frontal cortex of the rat: dopaminergic innervation of the prefrontal subareas and lateralization of prefrontal dopamine. Brain Res. 1982 Oct 28;250(1):197–200.

24. Adams RN. Probing brain chemistry with electroanalytical techniques. Anal Chem. 1976;48(14):1126A-1138A.

25. Michael DJ, Wightman RM. Electrochemical monitoring of biogenic amine neurotransmission in real time. J Pharm Biomed Anal. 1999;19(1):33–46.

26. Shnitko TA, Robinson DL. Anatomical and pharmacological characterization of catecholamine transients in the medial prefrontal cortex evoked by ventral tegmental area stimulation. Synap N Y N. 2014 Apr;68(4):131–43.

27. Van Der Weele CM, Siciliano CA, Matthews GA, Namburi P, Izadmehr EM, Espinel IC, et al. Dopamine enhances signal-to-noise ratio in cortical-brainstem encoding of aversive stimuli. Nature. 2018 Nov;563(7731):397–401.

28. Lohani S, Martig AK, Underhill SM, DeFrancesco A, Roberts MJ, Rinaman L, et al. Burst activation of dopamine neurons produces prolonged post-burst availability of actively released dopamine. Neuropsychopharmacology. 2018 Sep;43(10):2083–92.

29. Iwashita M. Phasic activation of ventral tegmental neurons increases response and pattern similarity in prefrontal cortex neurons. Häusser M, editor. eLife. 2014 Sep 30;3:e02726.

30. Kuznetsova A, Brockhoff PB, Christensen RHB. lmerTest Package: Tests in Linear Mixed Effects Models. J Stat Softw. 2017 Dec 6;82(13):1–26.

31. Huys QJM, Cools R, Gölzer M, Friedel E, Heinz A, Dolan RJ, et al. Disentangling the Roles of Approach, Activation and Valence in Instrumental and Pavlovian Responding. PLOS Comput Biol. 2011 Apr 21;7(4):e1002028.

### Supplementary Figures and Legends

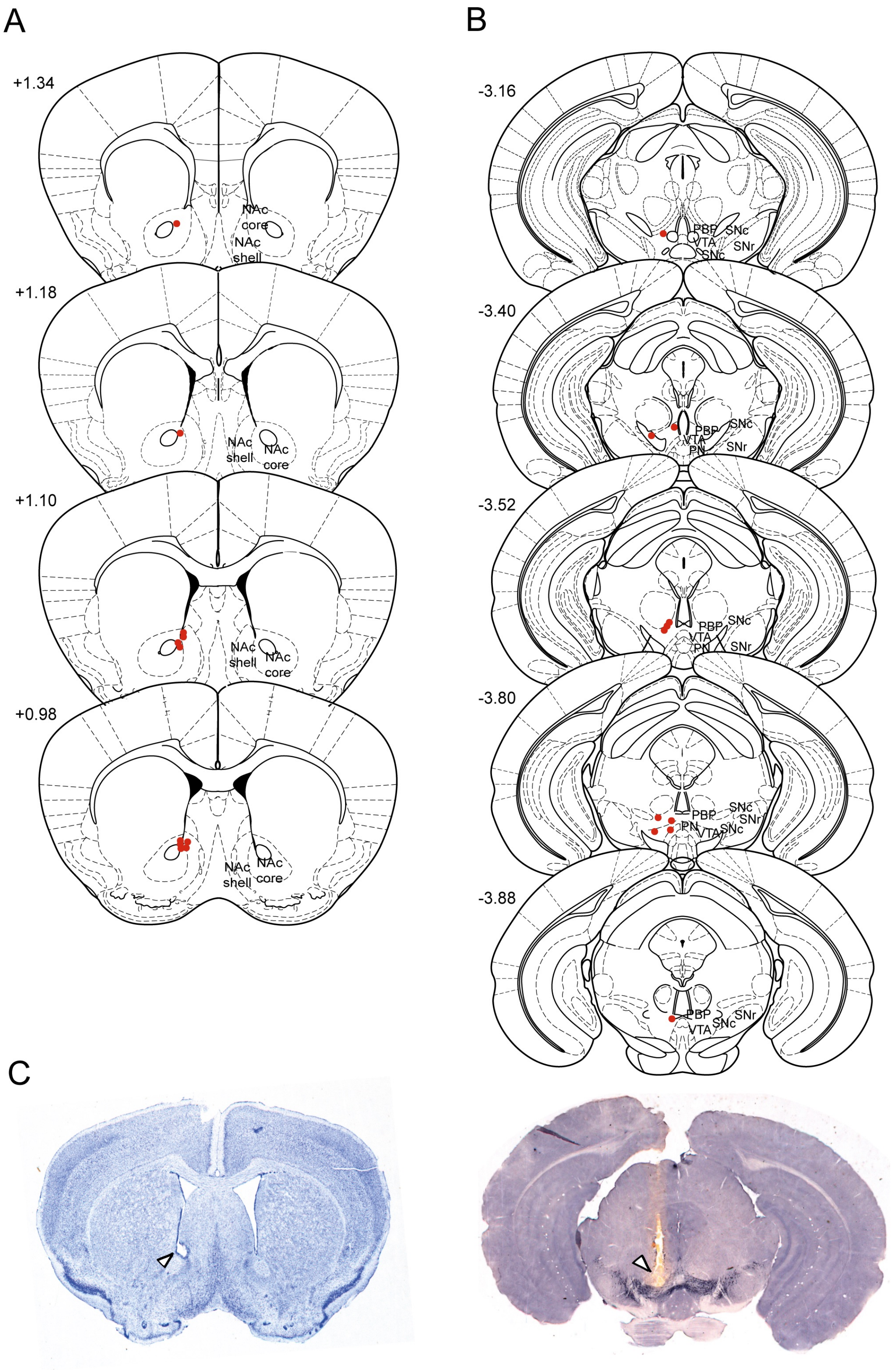

**Supplementary Figure 1: Electrode placements for NAc voltammetry recordings**

**A)** Locations of electrolytic lesions made at recording electrode sites in the subset of animals in which lesions were made (11 out of 23 total animals). **B)** Locations of the ends of stimulating electrode tracts. **C)** Examples of a recording electrode lesion in the NAc (left) and a stimulating electrode tract terminating in the VTA (right), indicated by white arrowheads. The VTA histological section was stained for tyrosine hydroxylase and demonstrates the proximity of the stimulating electrode tract to dopaminergic cell bodies (black) in the midbrain. Numbers to the left of coronal sections in (A) and (B) indicate distance from bregma (mm). NAc: nucleus accumbens; PBP: parabrachial pigmented nucleus; PN: paranigral nucleus; SNc: substantia nigra pars compacta; SNr: substantia nigra pars reticulata; VTA: ventral tegmental area.

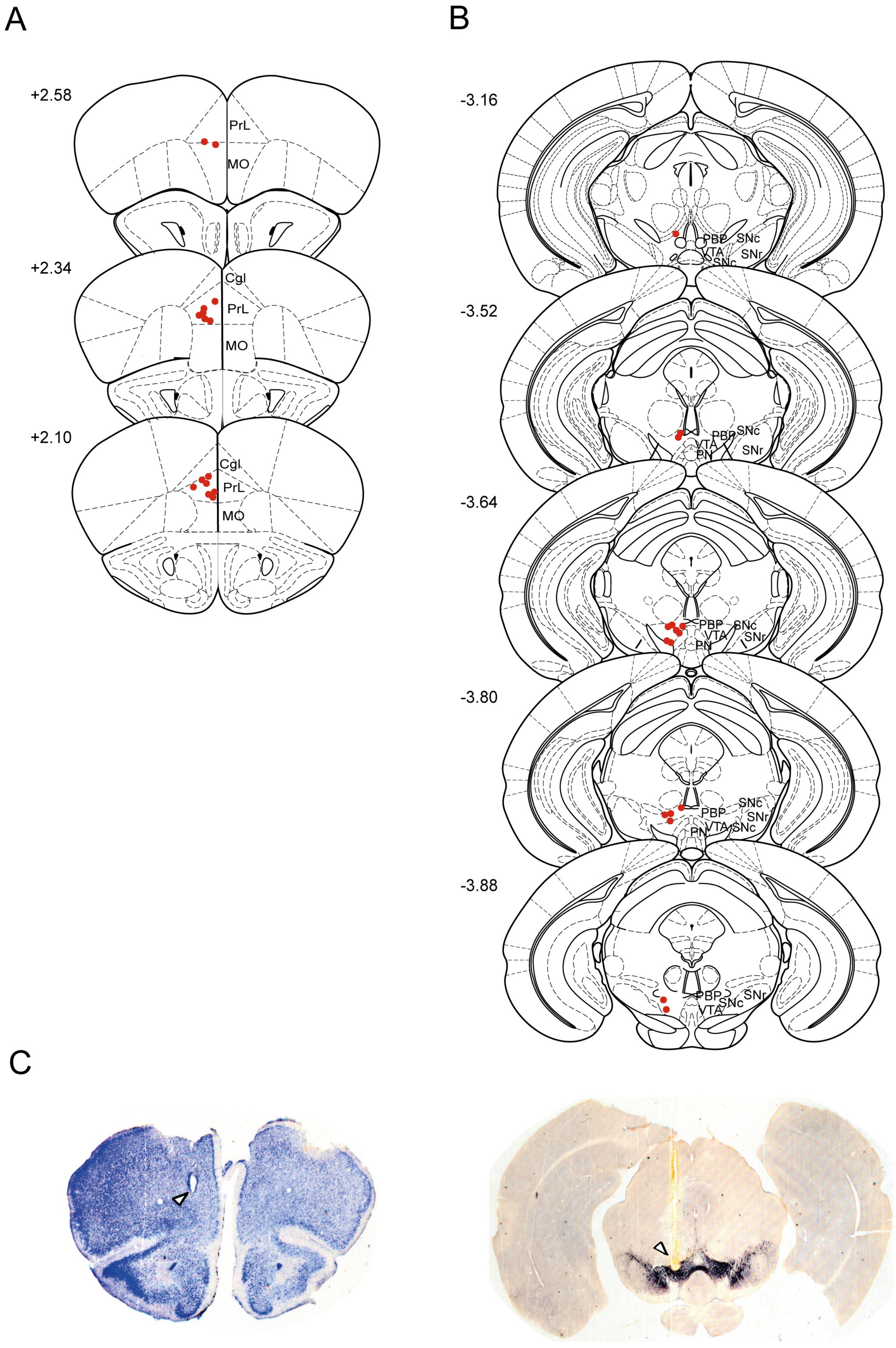
A
B
C
+2.58
PrL
MO
+2.34
Cgl
PrL
MO
+2.10
Cgl
PrL
MO
-3.16
PBP
SNc
VTA
SNr
SNc
-3.52
PBP
SNc
VTA
PN
SNr
-3.64
PBP
SNc
VTA
SNr
PN
-3.80
SNc
PBP
SNr
VTA
SNc
PN
-3.88
PBP
SNc
SNr
VTA

**Supplementary Figure 2: Electrode placements for PFC voltammetry recordings**

**A)** Locations of electrolytic lesions made at recording electrode sites in the subset of animals in which lesions were made (16 out of 34 total animals). **B)** Locations of the ends of stimulating electrode tracts. **C)** Examples of a recording electrode lesion in the PFC (left) and a stimulating electrode tract terminating in the VTA (right), indicated by white arrowheads. The VTA histological section was stained for tyrosine hydroxylase and demonstrates the proximity of the stimulating electrode tract to dopaminergic cell bodies (black) in the midbrain. Numbers to the left of coronal sections in (A) and (B) indicate distance from bregma (mm). Cgl: cingulated cortex; MO: medial orbital cortex; PBP: parabrachial pigmented nucleus; PN: paranigral nucleus; PrL: prelimbic cortex; SNc: substantia nigra pars compacta; SNr: substantia nigra pars reticulata; VTA: ventral tegmental area.

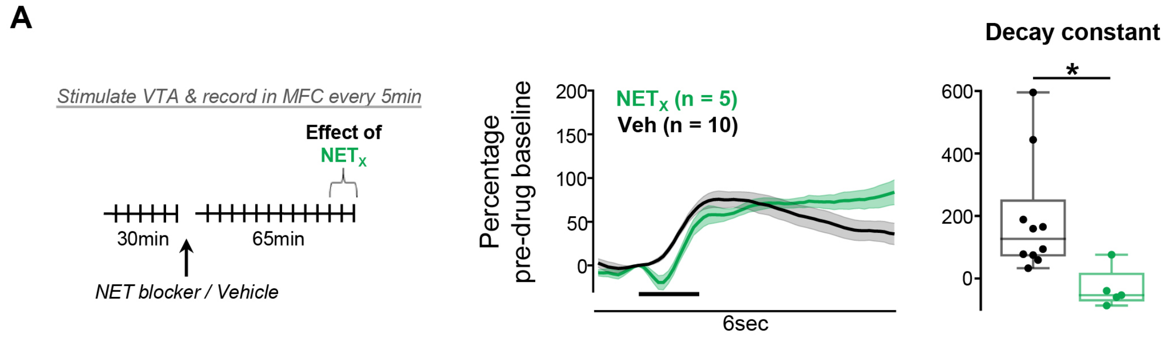

**Supplementary Figure 3: Effects of NET blockade on evoked dopamine release in the PFC**

**A)** The effect of the NET blocker atomoxetine (green) on evoked dopamine release in PFC. Left: schematic of experiment structure. Middle: evoked dopamine release normalized to the average pre-drug baseline peak height, binned over 15min centered on the time point of interest (60min after drug injection), and presented as mean ± SEM across animals within each group. Right: quantification of the signal decay, with each animal's data shown individually and normalized to its average pre-drug baseline decay constant. **Statistical analysis:** Atomoxetine slowed clearance of cortical transients compared to vehicle controls (independent samples t-test: $t_{13}$=2.6, p=0.023) but did not affect transient peak height ($t_{13}$=0.1, p=0.915).

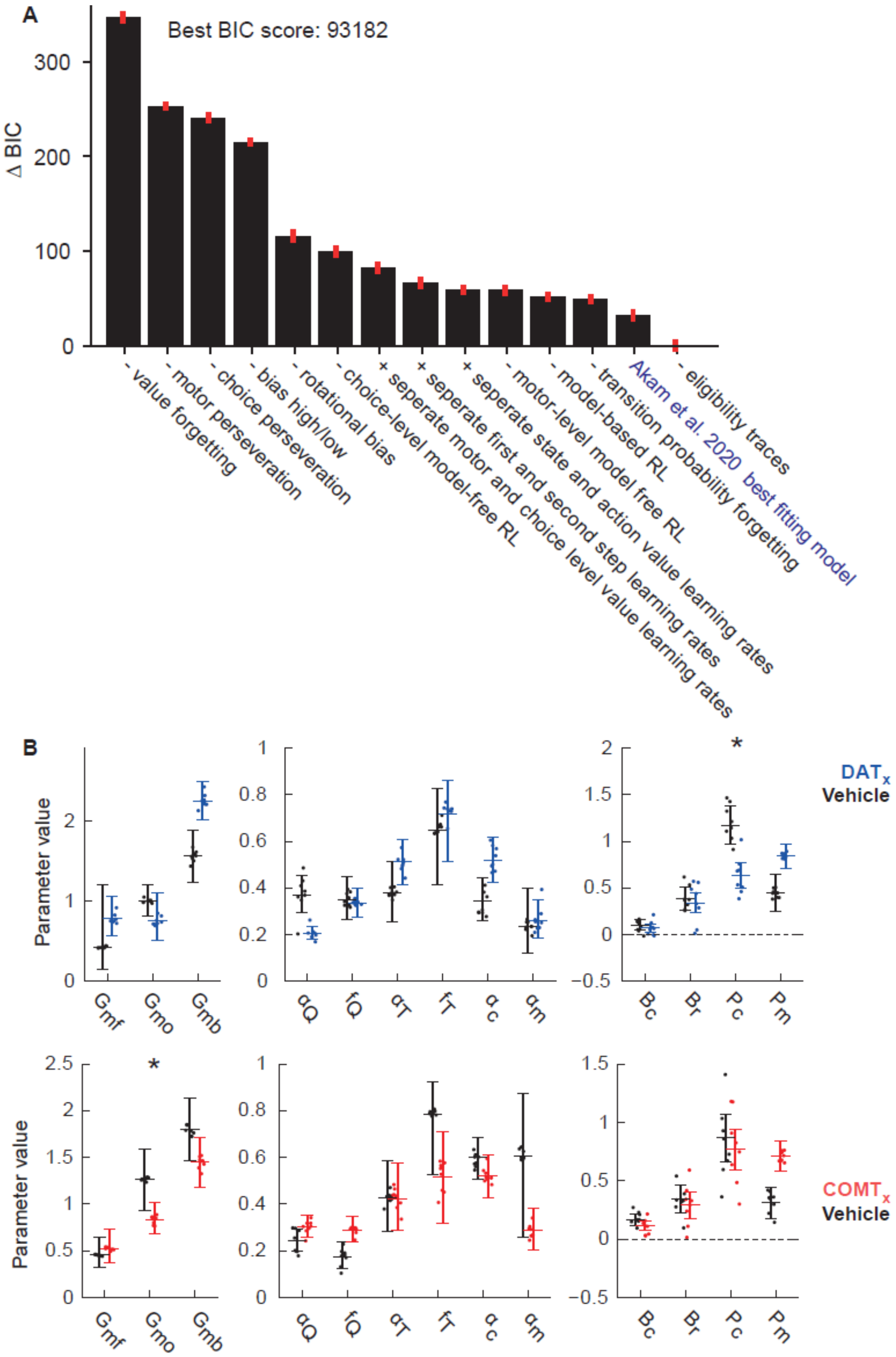
A
Best BIC score: 93182
Δ BIC
- value forgetting
- motor perseveration
- choice perseveration
- bias high/low
- rotational bias
- choice-level model-free RL
+ seperate motor and choice level value learning rates
+ seperate first and second step learning rates
+ seperate state and action value learning rates
- motor-level model free RL
- model-based RL
- transition probability forgetting
Akam et al. 2020 best fitting model
- eligibility traces
B
Parameter value
Gmf
Gmo
Gmb
αQ
fQ
αT
fT
αc
αm
Bc
Br
Pc
Pm
DATx
Vehicle
COMTx
Vehicle

**Supplementary Figure 4: Reinforcement learning modelling of multi-step decision making behavior A)** Model comparison showing BIC score differences, relative to the best fitting model, for RL agents fitted to choices on baseline testing days (days with neither drug nor vehicle injections). The set of models considered were generated by adding or removing single features from the model found to best describe behavior on this task in (18). Models are labelled on the x-axis by the feature that was added or removed. **B)** Model fits to data from drug and vehicle sessions for DAT blockade (top panel) and COMT inhibition (bottom panel) manipulations. Dots show maximum a posteriori fits for individual subjects, bars show 95% confidence intervals on the population means estimated using the Hessian of the log likelihood.

### Supplementary tables

**Supplementary Table 1:** Voltammetry experiment group sizes.

| Region & experiment | Group size for each drug treatment | |
|---|---|---|
| *NAc COMT inhibition* | Vehicle: n = 12<br>(n = 5 Veh/Veh, n = 7 Veh/GBR) | Tolcapone: n = 11<br>(n = 6 Tolc/Veh, n = 5 Tolc/GBR) |
| *NAc DAT blockade* | Vehicle: n = 11<br>(n = 5 Veh/Veh, n = 6 Tolc/Veh) | GBR-12909: n = 12<br>(n = 7 Veh/GBR, n = 5 Tolc/GBR |
| *PFC COMT inhibition* | Vehicle: n = 11<br>(n = 6 Veh/Veh, n = 5 Veh/GBR) | Tolcapone: n = 14<br>(n = 6 Tolc/Veh, n = 8 Tolc/GBR) |
| *PFC DAT blockade* | Vehicle: n = 12<br>(n = 6 Veh/Veh, n = 6 Tolc/Veh) | GBR-12909: n = 13<br>(n = 5 Veh/GBR, n = 8 Tolc/GBR) |
| *PFC amphetamine* | Amphetamine: n = 7<br>(n = 1 Veh/Veh/Amph, n = 2 Tolc/Veh/Amph, n = 4 Amph only)<br>Vehicle: n = 10<br>(n = 5 Veh/Veh, n = 5 Veh/GBR, at timepoint just prior to administration of GBR. NB. Veh/Veh animals that later received Amph were not included in the control group) | |
| *PFC atomoxetine* | Atomoxetine: n = 5<br>Vehicle: n = 10<br>(see *PFC amphetamine* for details) | |

**Supplementary Table 2:** Multi-step decision making task parameter changes over training.

| Session number | Reward size (µl) | Transition probabilities (common / rare) | Reward probabilities (good / bad side) |
|---|---|---|---|
| 1 | 4 or 8 | 0.9 / 0.1 | First 40 trials all rewarded, subsequently 0.9 / 0.1 |
| 2 - 18 | 4 or 8 | 0.9 / 0.1 | 0.9 / 0.1 |
| 19 - 21 | 8 | 0.8 / 0.2 | 0.9 / 0.1 |
| 22 - 23 | 6.5 | 0.8 / 0.2 | 0.9 / 0.1 |
| 24 - 26 | 6.5 | 0.8 / 0.2 | 0.8 / 0.2 |
| 27+ | 4 | 0.8 / 0.2 | 0.8 / 0.2 |

**Supplementary Table 3:** DAT blockade reversal analysis fit for the multi-step decision making task. Parameters of double exponential fit to reversals in reward and transition probabilities under GBR-12909 and vehicle.

| **Reversal type** | **Condition** | **Parameter** | | |
|---|---|---|---|---|
| | | $tau_F$ | $tau_S$ | $w_F$ |
| Reward | *Vehicle* | 2.58 | 280.12 | 0.70 |
| | *GBR-12909* | 15.68 | 7842.5 | 0.89 |
| | *p value* | 0.0012 | 0.0008 | 0.98 |
| Transition | *Vehicle* | 0.83 | 35.95 | 0.44 |
| | *GBR-12909* | 3.58 | 62.27 | 0.64 |
| | *p value* | 0.23 | 0.69 | 0.44 |

**Supplementary Table 4:** COMT inhibition reversal analysis fit for the multi-step decision making task. Parameters of double exponential fit to reversals in reward and transition probabilities under tolcapone and vehicle.

| **Reversal type** | **Condition** | **Parameter** | | |
|---|---|---|---|---|
| | | $tau_F$ | $tau_S$ | $w_F$ |
| Reward | *Vehicle* | 13.6 | 54.24 | 0.57 |
| | *Tolcapone* | 3.98 | 39.27 | 0.48 |
| | *p value* | 0.0496 | 0.84 | 0.84 |
| Transition | *Vehicle* | 5.03 | 2514 | 0.90 |
| | *Tolcapone* | 4.42 | 2207 | 0.76 |
| | *p value* | 0.72 | 0.75 | 0.3 |

**Supplementary Table 5:** RL model parameters and variables

| **Model variables** | |
|---|---|
| $r$ | reward (0 or 1) |
| $c$ | choice taken at first step (top or bottom poke) |
| $c'$ | choice not taken at first step (top or bottom poke) |
| $s$ | Second-step state (left-active or right-active) |
| $s'$ | State not reached at second step (left-active or right-active) |
| $Q_{mf}(c)$ | Model-free action value for choice $c$ |
| $Q_{mo}(c,\, s_{t-1})$ | Motor-level model-free action value for choice $c$ following second-step state $s_{t-1}$ |
| $Q_{mb}(c)$ | Model-based value of choice $c$ |
| $V(s)$ | Value of state $s$ |
| $P(s\|c)$ | Estimated transition probability of reaching state $s$ after choice $c$ |
| $\bar{c}$ | Choice history |
| $\bar{m}(s_{t-1})$ | Motor action history, i.e. choice history following second-step state $s_{t-1}$ |
| **Model parameters** | |
| $G_{mf}$ | Model-free action value weight |
| $G_{mo}$ | Motor-level model free action value weight |
| $G_{mb}$ | Model-based action value weight |
| $\alpha_Q$ | Value learning rate |
| $f_Q$ | Value forgetting rate |
| $\alpha_T$ | Transition learning rate |
| $f_T$ | Transition forgetting rate |
| $\alpha_c$ | Learning rate for choice perseveration |
| $\alpha_m$ | Learning rate for motor-level perseveration |
| $B_c$ | Choice bias (top/bottom) |
| $B_r$ | Rotational bias (clockwise/counter-clockwise) |
| $P_c$ | Choice perseveration strength |
| $P_m$ | Motor-level perseveration strength |

**Supplementary Table 6:** P value for differences in RL model parameter fits between drug and corresponding vehicle sessions for DAT blockade and COMT inhibition. P values <0.1 are indicated in blue, <0.05 in red. P values are not reported multiple comparison corrected for the 13 parameters in the model.

| Parameter | Description | DATx p value | COMTx p value |
|---|---|---|---|
| $G_{mf}$ | Model-free action value weight | 0.094 | 0.814 |
| $G_{mo}$ | Motor-level model free action value weight | 0.368 | 0.028 |
| $G_{mb}$ | Model-based action value weight | 0.125 | 0.629 |
| $\alpha_Q$ | Value learning rate | 0.073 | 0.410 |
| $f_Q$ | Value forgetting rate | 0.736 | 0.066 |
| $\alpha_T$ | Transition learning rate | 0.246 | 0.860 |
| $f_T$ | Transition forgetting rate | 0.464 | 0.152 |
| $\alpha_c$ | Learning rate for choice perseveration | 0.068 | 0.486 |
| $\alpha_m$ | Learning rate for motor-level perseveration | 0.923 | 0.055 |
| $B_c$ | Choice bias (top/bottom) | 0.424 | 0.078 |
| $B_r$ | Rotational bias (clockwise/counter-clockwise) | 0.490 | 0.451 |
| $P_c$ | Choice perseveration strength | 0.027 | 0.987 |
| $P_m$ | Motor-level perseveration strength | 0.130 | 0.148 |
